# High-sensitivity monitoring of ctDNA by patient-specific sequencing panels and integration of variant reads

**DOI:** 10.1101/759399

**Authors:** Jonathan C. M. Wan, Katrin Heider, Davina Gale, Suzanne Murphy, Eyal Fisher, James Morris, Florent Mouliere, Dineika Chandrananda, Andrea Marshall, Andrew B. Gill, Pui Ying Chan, Emily Barker, Gemma Young, Wendy N. Cooper, Irena Hudecova, Francesco Marass, Graham R. Bignell, Constantine Alifrangis, Mark R. Middleton, Ferdia A. Gallagher, Christine Parkinson, Amer Durrani, Ultan McDermott, Christopher G. Smith, Charles Massie, Pippa G. Corrie, Nitzan Rosenfeld

**Author notes:** J.C.M.W. and K.H. contributed equally to this work. C.M., P.G.C. and N.R. jointly supervised this work. **One Sentence Summary:** Integrating tumor-derived sequences across large panels of patient-specific mutations offers enhanced sensitivity for ctDNA detection and monitoring from both high-depth and low-depth plasma sequencing data.

## Abstract

Circulating tumor-derived DNA (ctDNA) can be used to monitor cancer dynamics noninvasively. Patients with small tumors have few copies of ctDNA in plasma, resulting in limited sensitivity to detect low-volume or residual disease. We show that sampling limitations can be overcome and sensitivity for ctDNA detection can be improved by massively parallel sequencing when hundreds to thousands of mutations are identified by tumor genotyping. We describe the INtegration of VAriant Reads (INVAR) analysis pipeline, which combines patient-specific mutation lists with both custom error-suppression methods and signal enrichment based on biological features of ctDNA. In this framework, the sensitivity can be estimated independently for each sample based on the number of informative reads, which is the product of the number of mutations analyzed and the average depth of unique sequencing reads. We applied INVAR to deep sequencing data generated by custom hybrid-capture panels, and showed that when ~10^6^ informative reads were obtained INVAR allowed detection of tumor-derived DNA fractions to parts per million (ppm). In serial samples from patients with advanced melanoma on treatment, we detected ctDNA when imaging confirmed tumor volume of ~1cm^3^. In patients with resected early-stage melanoma, ctDNA was detected in 40% of patients who later relapsed, with higher rates of detection when more informative reads were obtained. We further demonstrated that INVAR can be generalized and allows improved detection of ctDNA from whole-exome and low-depth whole-genome sequencing data.

## Introduction

Circulating tumor DNA (ctDNA) can be robustly detected in plasma when multiple copies of mutant DNA are present; however, when ctDNA levels are low, analysis of individual mutant loci might produce a negative result due to sampling noise even when using an assay with perfect analytical sensitivity. Such “missed” samples can have low fractional concentrations of ctDNA (relatively few mutant molecules in a high background), or low absolute numbers of mutant molecules due to limited sample input (Fig. 1A). This effect of limited sampling reduces the sensitivity of ctDNA monitoring for patients with early-stage cancers, or following treatment for detection of minimal residual disease *(1*, *2)*. Studies showed, for example, that by targeting a single mutation per patient in the plasma of early-stage breast and colorectal cancer patients post-operatively, ctDNA was detected in approximately 50% of patients who later relapsed *(3*, *4)*. When applied to *BRAF*- or *NRAS*-mutant stage II-III patients with melanoma, ctDNA was detected up to 12 weeks post-surgery in only 16.8% of patients who relapsed within 5 years *(5)*. To increase the number of mutant molecules sampled, previous studies have shown that it may be possible to analyze larger volumes of plasma from multiple blood tubes *(4*, *6)* and/or utilize broader sequencing panels.

**Fig. 1.**
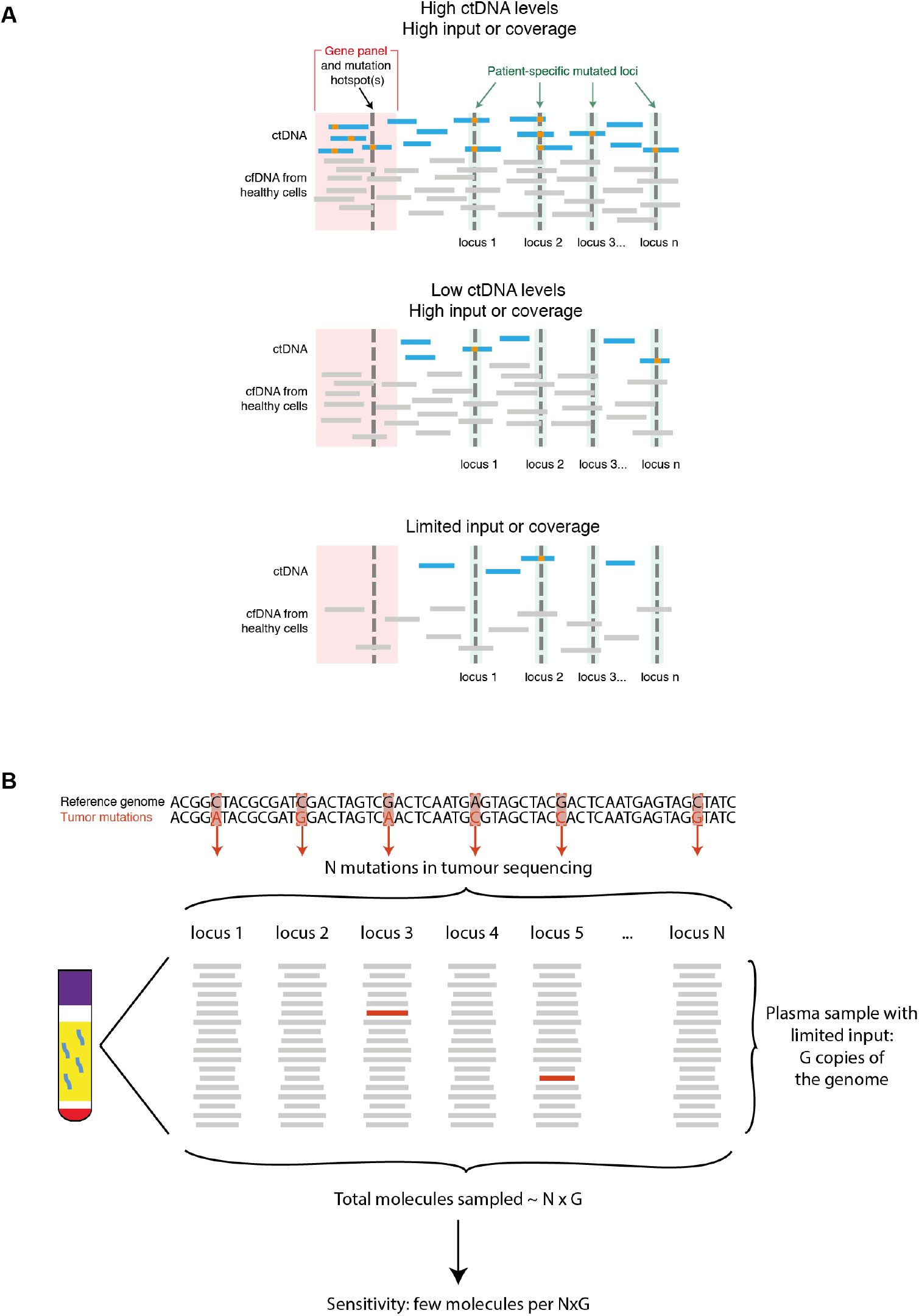
Patient-specific analysis overcomes sampling error in conventional and limited input scenarios. (**A**) When high levels of ctDNA are present, gene panels and hotspot analysis are sufficient to detect ctDNA (top panel). However, if ctDNA concentrations are low these assays are at high risk of false negative results due to sampling noise. Utilizing a large list of patient specific mutations allows sampling of mutant reads at multiple loci, enabling detection of ctDNA when there are few mutant reads due to either ultra-low ctDNA levels (middle panel), or due to limited starting material or sequencing coverage (bottom panel). (**B**) A given sample contains a limited number of copies of the genome, denoted by G. For plasma samples, the small amount of material limits the sensitivity that is attainable to one mutant copy in G total copies. By analyzing in parallel a large number of marker loci (e.g. loci that are found to be mutated in the patient’s tumor), denoted by N, detection of tumor DNA can be substantially enhanced to detect one or few mutant molecules per N × G copies. The same approach can be employed for other applications which aim to detect non-background/altered DNA, such as detection of fetal DNA or DNA from transplanted organs, in limited amounts of material such as plasma samples or other body fluids.

Tumor-guided patient-specific analysis, which involves prior tumor genotype information and custom panel design *(7*–*11)*, offers the possibility to greatly increase the sensitivity of ctDNA assays for cancer monitoring by targeting a larger number of mutations *(2*, *11)* (Fig. 1B). Such assays have analyzed up to 40 patient-specific mutations in parallel, quantifying ctDNA to 1 mutant molecule per 25,000 copies in a patient with non-small cell lung cancer (NSCLC) *(10)*. Increasingly broad tumor sequencing is being performed both in research and clinical settings *(12)*, which provides valuable mutation information that may be leveraged for improved sensitivity for ctDNA. We conceptualize the factors influencing ctDNA sensitivity as a two-dimensional space (Fig. 2A), highlighting the importance of maximizing the number of relevant DNA fragments analyzed, by increasing either plasma volumes or the number of (patient-specific) mutations sampled: the number of informative reads (IR) generated is proportional to the product of these two factors.

**Fig. 2.**
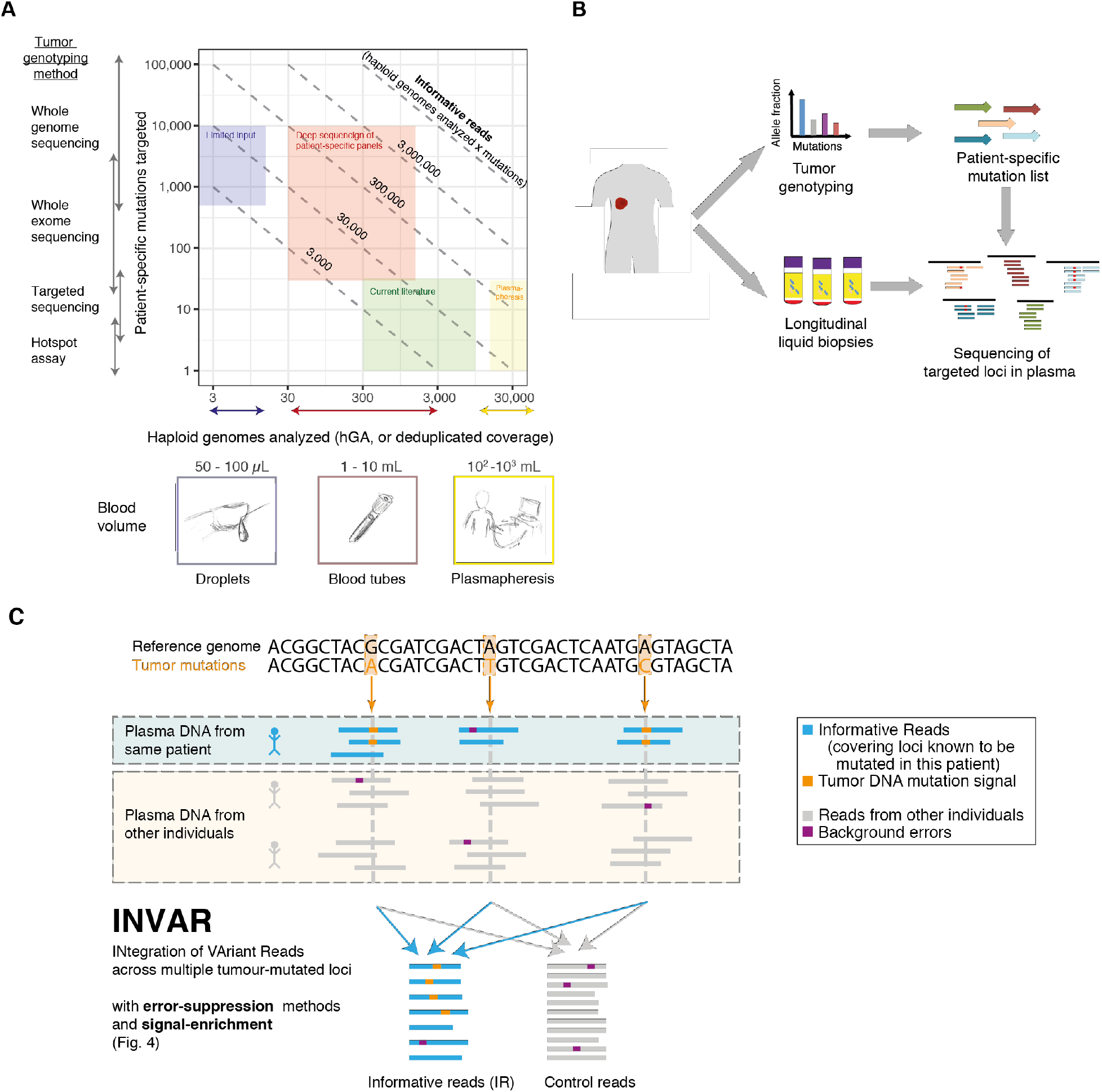
Study outline and rationale for integration of variant reads. (**A**) Illustration of the range of possible working points for ctDNA analysis using INVAR, plotting the haploid genomes analyzed vs. the number of mutations. Diagonal lines indicate multiple ways to generate the same number of informative reads (IR, equivalent to haploid genomes analyzed (hGA) x targeted loci). Current methods often focus on analysis of ~10 ng of DNA (300-10,000 haploid copies of the genome) across 1 to 30 mutations per patient. This typically results in ~10,000 IR, leading to frequently encountered detection limits of 0.01%-0.1% *(9*, *13)*. In this study we focused on analysis of larger numbers (100s-1000s) of mutations, including deep sequencing by patient-specific hybrid-capture panels or limited input. mL, milliliter; μL, microliter. (**B**) To generate deep sequencing data across large patient-specific mutations lists at high depth, patient-specific mutation lists generated by tumor genotyping were used to design hybrid-capture panels, that were applied to DNA extracted from plasma samples. In later sections, the tumor genotyping data is used to analyze sequencing data from standard WES panels and shallow WGS. (**C**) The INtegration of VAriant Reads (INVAR) pipeline. To overcome sampling error, signal was aggregated across hundreds to thousands of mutations. Here we classify samples (rather than individual mutations) as significantly containing ctDNA, or not detected. ‘Informative Reads’ (IR, shown in blue) are reads generated from a patient’s sample that overlap loci in the same patient’s mutation list. Some of these reads may carry the mutation variants in the loci of interest (shown in orange). Reads from plasma samples of other patients at the same loci (‘non-patient-specific’) are used as control data to calculate the rates of background error rates (shown in purple) that can occur due to sequencing errors, PCR artefacts, or biological background signal. INVAR incorporates additional sequencing information on fragment length and tumor allelic fraction to enhance detection.

ctDNA detection methods often rely on identification of individual mutations *(6*, *9*, *13)* which may discard mutant signal that does not pass a threshold for calling. In this study, to improve sensitivity, we aggregated sequencing reads across 10^2^−10^4^ mutated loci, using prior information from tumor genotyping to guide analysis (Fig. 2B). The potential sensitivity benefit of targeting hundreds to thousands of tumor markers per patient has been previously suggested *(10*, *14)*, though such approaches have not been applied to cancer monitoring in plasma.

We suggest that a tumor-guided approach targeting a large number of patient-specific mutations has advantages beyond simply mitigating sampling error. By virtue of generating a large number of IR, multiple error-suppression steps may be employed to overcome sequencing and PCR errors while retaining signal. Aside from molecular barcoding, it may be possible to identify artefactual signal at a given locus by comparison of the given allelic fraction against the allele fractions at other patient-specific loci. Furthermore, greater weight may be assigned to fragments more likely to arise from tumor cells based on their biological characteristics such as fragment size *(15)*, thereby enhancing the signal to noise ratio.

Here, we present a workflow for enhanced patient-specific monitoring that is optimized for sensitive detection of ctDNA to parts per million, using patient-specific sequencing data and custom hybrid-capture panels (Fig. 2C, flowchart in Fig. S1). This approach leverages custom error-suppression and signal enrichment methods to enable sensitive monitoring and identification of residual disease. We further demonstrate the ability to apply INVAR to plasma whole-exome sequencing (WES) and shallow whole genome sequencing (sWGS), demonstrating improved sensitivity for detection and quantification of ctDNA.

### Tumor genotyping

First, tumor genotyping was performed to identify multiple patient-specific mutations per patient: exome sequencing data was generated from tumor and buffy coat samples from 47 patients with Stage II-IV melanoma (Methods), identifying a median of 625 mutations per patient (IQR 411-1076, Fig. S2 and Table S1). These mutation lists were used to generate custom capture sequencing panels, which were used to sequence longitudinal plasma samples (n=144) (2,301x mean raw depth). In addition, WES (238x mean raw depth, n=20) and sWGS (0.6x mean raw depth, n=33), was performed on samples from the same patients and used as input for INVAR analysis (Tables S2 and S3).

### Characterizing background error rates

We started by characterizing background error rates in hybrid-capture sequencing data. Approximation of error rates may potentially be achieved through grouping mutations of similar class. We demonstrate that error rates vary between mutation class by over an order of magnitude using raw sequencing data without using molecular barcodes (Fig. 3A), consistent with Newman et al. *(10)*. To increase the resolution of background error rates further, we grouped mutations by both mutation class and trinucleotide context, demonstrating over two orders of magnitude difference in background error rate between the least and most noisy trinucleotide context (Fig. 3B).

**Fig. 3.**
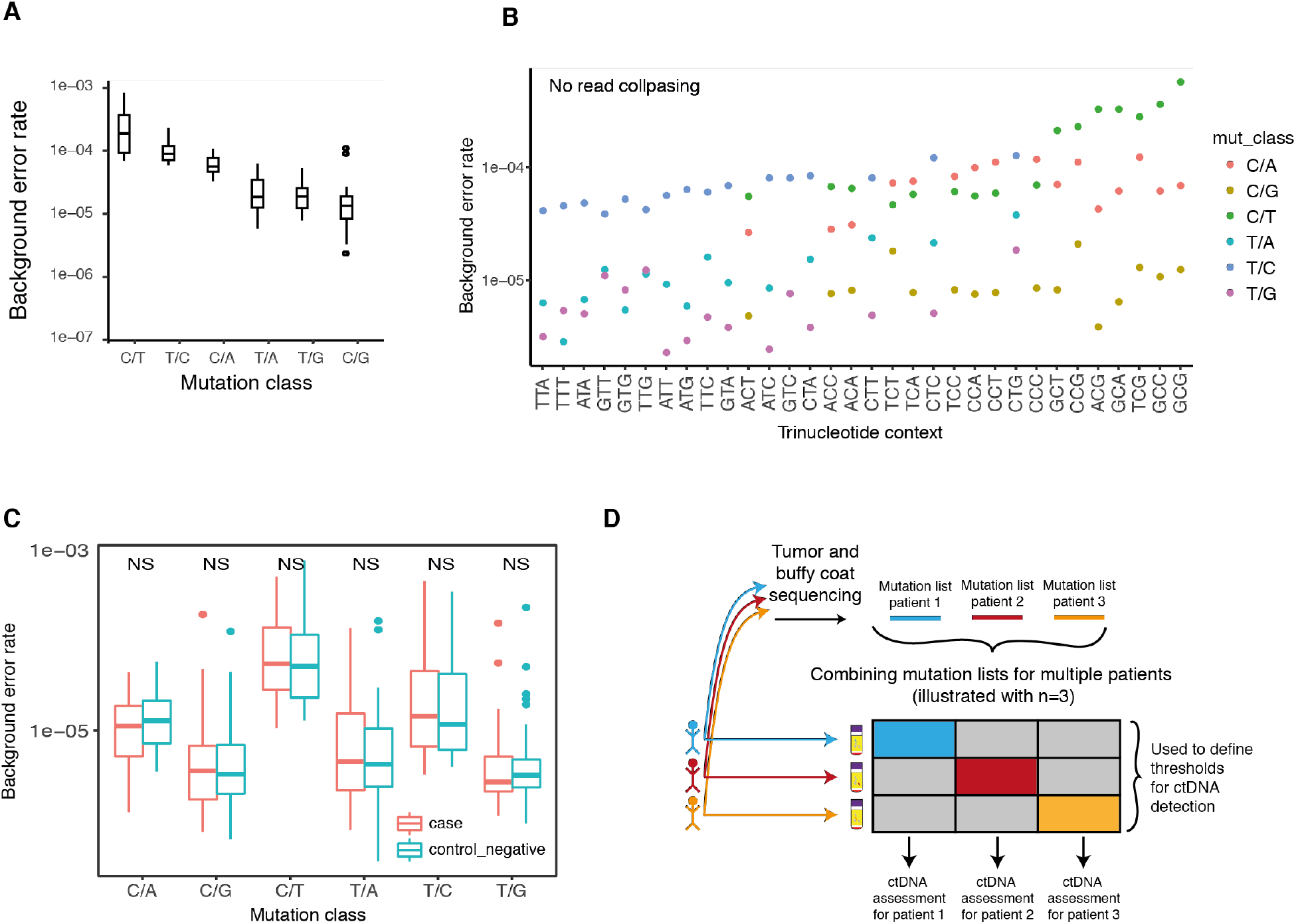
Characterization of background error rates. (**A**) Non-error suppressed background error rates by mutation class. (**B**) Non-error suppressed background error rates by trinucleotide context. (**C**) Background error rates were calculated by mutation class for healthy control individuals (blue) and patient samples (red) after equalizing the number of read families per group. Complementary mutation classes were combined. T-tests were performed between healthy and patient samples. NS, not significant. (**D**) Overview of the usage of sequencing data by the INVAR pipeline. Data is collected for each locus of interest in the matched patient (shown by colored boxes), and control data is obtained by analyzing the same loci in additional patients from the same cohort for whom the loci were not found to be mutated in the tumor or buffy coat analysis. Such data can be generated by applying a standardized sequencing approach, such as WES/sWGS, to all samples (Fig. 7) or by combining multiple patient-specific mutation lists into a custom capture panel that is sequenced across multiple patients (Fig. 6). Data from other patients in those loci (‘non-matched mutations’) are used to determine background mutation rates and a ctDNA detection cut-off.

Using a patient-specific sequencing approach, a large number of private mutation loci were targeted. Each locus has its own error rate, though accurate benchmarking of the background noise rates of individual loci to levels below 10^−6^ would require the cfDNA molecules from a total of 100mL of plasma in order to sample one mutant read. This assumes a cfDNA concentration of 10ng/mL from plasma, yielding 3 million analyzable molecules. Thus, we sought to develop a background error model for patient-specific sequencing data that could estimate the background error rate of a locus accurately using limited control samples. In this study, 99.8% of the mutations identified by tumor sequencing were private i.e. unique to each individual. We assessed if patients may be used to control for other patients’ mutation lists, thereby enabling us to group patient-specific mutation lists of multiple patients together and reduce the number of additional control samples to be run on each panel. In this study, a mean of 5.5 patients were included on each custom hybrid-capture sequencing panel design. There was no significant difference in background error rate whether using healthy individuals or other patients serving as controls (‘patient-control’ samples, which may control for other patients at private loci) (Fig. 3C). Thus, INVAR utilizes sequencing data from one patient to control for others with both custom and untargeted approaches such as exome or WGS (Fig. 3D).

### Error-suppression in patient-specific sequencing data

As part of the INVAR pipeline, we sought to develop methods to minimize artefacts in patient-specific sequencing data. Read collapsing was performed using unique molecular barcodes which reduced error rates across all mutation classes (Fig. S3A), similar to previous studies *(16)*. Increasing the minimum number of duplicates required per read family reduced error rates further, but at the expense of a greater fraction of the sequencing data being discarded (Fig. S3B). To balance data loss against background error rate, a minimum family size threshold of 2 was used.

INVAR requires any mutation signal to be represented in both the forward (F) and reverse (R) read of the read pair. This serves to both reduce sequencing error and produces a small size-selection effect for short fragments since only short fragments would be read completely in both F and R with paired-end 150bp sequencing. This step retained 92.4% of mutant reads and 84.0% of wild-type reads in a training dataset (Fig. S3C).

When targeting a large number of patient-specific loci, it becomes increasingly likely that technically noisy sites, or single-nucleotide polymorphism (SNP) loci are included in the list. Newman et al. have previously utilized position specific polishing to address this issue *(10)*. In this study we blacklisted loci that showed either error-suppressed mutant signal in >10% of the patient-control samples, or a mean background error rate of >1% mutant allele fraction. This approach excluded 0.5% of the patient-specific loci (Fig. S3D). Requiring mutant signal in both reads and applying a locus noise filter reduced noise modestly when applied individually; however, when combined they showed a synergistic effect, reducing background error rates to below 1×10^−6^ in some mutation classes (Figs. 4A, S3E). The individual effects of these filters on individual trinucleotide contexts are shown in Fig. S3F.

**Fig. 4.**
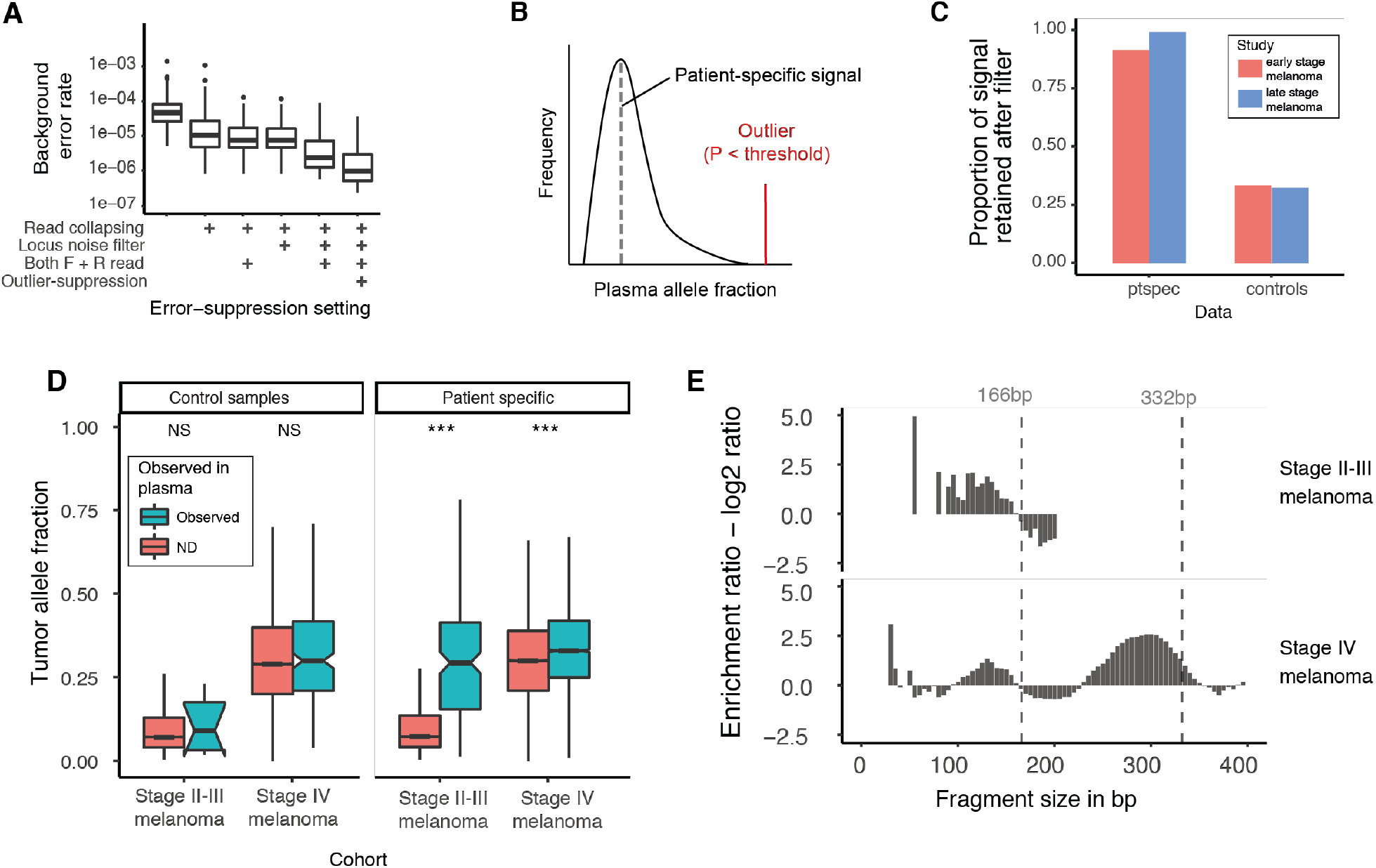
Development and analytical performance of the INVAR method. (**A**) Reduction of error rates following different error-suppression settings (Supplementary Methods). (**B**) Loci observed with significantly greater signal than the remainder of the loci of that patient might be due to noise at that locus, contamination, or a mis-genotyped SNP locus (in red, see Methods). (**C**) Summary of effect of outlier suppression on both cohorts. Mutant signal was reduced 3-fold in control samples, while retaining 96.1% of mutant signal in patient samples. (**D**) Tumor allele fractions were compared between loci with and without detected signal in plasma. Loci with signal in plasma had significantly higher tumor allele fractions in patient samples. There was no significant increase in tumor allele fraction when performing this analysis on non-patient-specific samples (Student’s t-test, NS, not significant; *** = P < 0.0001). (**E**) Log_2_ enrichment ratios for mutant fragments from three different cohorts of patients. Size ranges enriched for ctDNA are assigned more weight by the INVAR pipeline.

When targeting a large number of patient-specific sites, it becomes possible to assess the distribution of allele fractions observed. In the residual disease setting, we expect to have a high degree of sampling error. Therefore, signal should appear stochastically as individual mutant molecules distributed across patient-specific loci, with many of the loci having zero mutant reads. In order to optimize INVAR for detection of the lowest possible levels of ctDNA, we developed a method called patient-specific outlier suppression to exclude signal at a locus that is not consistent with the remaining loci (Figs. S3G and 4B). This tests each locus against the distribution of signal at all other loci with a correction for multiple testing, excluding loci that are significantly outlying. Mutant signal was reduced 3-fold in control samples, while retaining 96.1% of mutant signal in patient samples (Fig. 4C).

Overall, combining the above steps results in an average 131-fold decrease in background error relative to raw sequencing data (Fig. 4A) and reduces the error rates of some trinucleotide contexts to below 10^−6^ (Fig. S3E).

### Patient-specific signal enrichment

To enhance detection further, INVAR is able to enrich for ctDNA signal through probability weighting based on the tumor allele fraction of each mutation locus and ctDNA fragment sizes. Tumor mutations with a higher tumor allelic fraction are more likely to be observed in the plasma *(8*, *17)*, therefore, greater weight was allocated to mutant signals in plasma from loci with high tumor mutant allele fraction. Using a dilution series, we confirm the relationship between the tumor allele fraction of a locus and rate of detection of ctDNA of that locus in plasma (Fig. S4A). We confirm in clinical samples that patient-specific mutation loci observed in plasma had a significantly higher tumor allele fraction compared to those not observed in plasma (P = 2×10^−16^; Wilcoxon test, Fig. 4D).

Analysis of 144 samples showed a nucleosomal pattern of cfDNA fragmentation, with mutant fragments shorter than wild-type fragments at the mono-nucleosome and di-nucleosome peak (Fig. S4B). We also observed that stage IV melanoma patients had a significantly higher median mutant fragment size compared to the stage II-III melanoma patients (163bp vs. 154bp, P = 2×10^−16^, Wilcoxon test, Fig. S4C). Previous research has shown enrichment for ctDNA when shorter fragments are selected using either in vitro or *in silico* size selection *(15*, *18*, *19)*. However, at low levels of signal, such methods can cause loss of rare mutant alleles *(20)*. Thus, in this study we weighted each signal based on its fragment size in order to boost ctDNA signal, while retaining all the data (Fig. 4E). Based on smoothed size profiles of mutant and wild-type fragments observed (Fig. S4D), patient data were used to size-weight other patients’ data using a leave-one-out approach (Supplementary Methods).

Following signal weighting, INVAR aggregates signal across all patient-specific mutations (Methods). In order to determine whether or not ctDNA is detected in a sample, data from non-matched mutations in other patients were used as negative controls to set the detection threshold (Fig. S5). An Integrated Mutant Allele Fraction (IMAF) is determined by taking a background-subtracted, depth-weighted mean allele fraction across the patient-specific loci in each sample (Supplementary Methods).

### Analytical sensitivity and specificity of INVAR

To benchmark the sensitivity of INVAR, we performed custom capture sequencing of a dilution series of plasma from one melanoma patient (stage IV disease), for whom we identified 5,073 mutations through exome sequencing. Plasma DNA from this patient was serially diluted into control volunteers’ plasma DNA to an expected IMAF of 3.6 × 10^−7^. Without use of unique molecular barcodes, INVAR detected ctDNA down to an expected allele fraction of 3.6 × 10^−5^, which was quantified to an average IMAF of 4.7 × 10^−5^ from two replicates (Fig. 5A). Following the use of molecular barcodes and custom error-suppression methods, the diluted ctDNA was detected to an expected IMAF of 3.6 × 10^−6^ (3.6 parts per million) in two replicates, with IMAF values of 4.3 and 5.2 ppm. Overall, the correlation between IMAF and the expected mutant fraction was 0.98 (Pearson’s r, p < 2.2 × 10^−16^, Fig. 5A). At an expected allele fraction of 3.6 × 10^−7^, ctDNA was detected in 2 out of 3 replicates. To assess the impact on sensitivity of the number of mutations targeted we downsampled sequencing data *in silico* to include subsets of patient-specific mutation lists. This confirmed that targeting more mutations resulted in more IR and correspondingly higher ctDNA detection rates (Fig. 5B, Supplementary Methods).

**Fig. 5.**
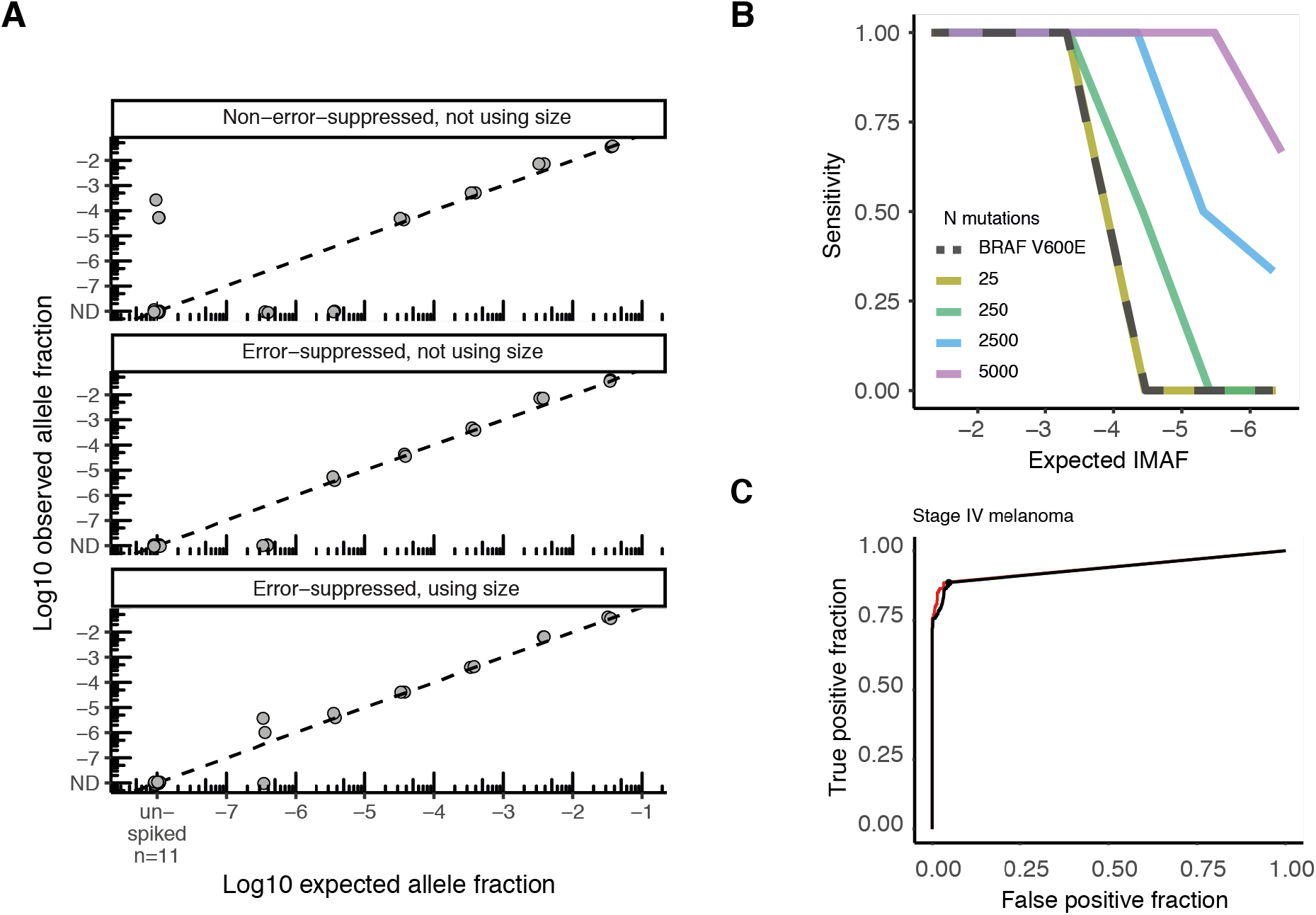
Sensitivity and specificity determination of INVAR. (**A**) Spike-in dilution experiment to assess the sensitivity of INVAR. Using error-suppressed data with INVAR, ctDNA was detected in replicates for all dilutions to 3.6 ppm, and in 2 of 3 replicates at an expected ctDNA allele fraction of 3.6 × 10^−7^ (Supplementary Methods). Using error-suppressed data of 11 replicates from the same healthy individuals without spiked-in DNA from the cancer patient, no mutant reads were observed in an aggregated 6.3 × 10^6^ informative reads across the patient-specific mutation list. (**B**) The sensitivity in the spike-in dilution series was assessed after the number of loci analyzed was downsampled *in silico* to between 1 and 5,000 mutations (Supplementary Methods). (**C**) ROC analysis of the stage IV melanoma cohort was performed against patient-controls (black) and healthy individuals (red).

The false positive rate of INVAR was measured twice, once in patient-control samples and separately in healthy control samples. First, analytical specificity was determined through analysis of samples from other patients (patient-control samples) at non-matched mutation loci, giving a median specificity of 98.0% (Fig. S6, Table S4). To confirm the specificity of INVAR in independent control samples, we ran custom capture sequencing (with the same oligo pools) on samples from healthy individuals and analyzed those by INVAR using each of the patient-specific mutation lists. The ROC curve for the stage IV melanoma cohort controlled against healthy individuals is shown in Fig. 5C. Across each of the analyses in this study, using control cfDNA from 26 healthy individuals, a median specificity value of 97.05% was obtained, consistent with the analytical specificity defined in non-matched control samples from other patients (Fig. S6).

### Quantification of ctDNA in patient samples

We applied INVAR to custom capture panel sequencing data from 130 plasma samples from 47 stage II-IV melanoma patients, generating up to 2.9 × 10^6^ IR per sample (median 1.7 × 10^5^ IR), thus analyzing orders of magnitude more cfDNA fragments compared to methods that analyze individual or few loci (Fig. 6A). In this study, we demonstrated a dynamic range of 5 orders of magnitude and detection of trace levels of ctDNA in plasma samples (Figs. 6B, 6C); this detection was obtained from a median input material of 1638 copies of the genome (5.46 ng of DNA; Table S2). In a total of 13 of the 130 plasma samples analyzed with custom capture sequencing, ctDNA was detected with signal in fewer than 1% of the patient-specific loci (Fig. 6D). The lowest fraction of cancer genomes detected was 1/714, equivalent to <5 femtograms of tumor DNA. Given the limited input, the low ctDNA levels detected would be below the 95% limit of detection for a ‘perfect’ single-locus assay in 48% of the cases (indicated with filled circles in Fig. 6C). The input mass vs. IMAF of each sample is shown in Fig. S7A, highlighting the sensitivity benefit of a broad sequencing approach. Thus, targeting multiple mutations can allow detection of low absolute amounts of tumor-derived DNA.

**Fig. 6.**
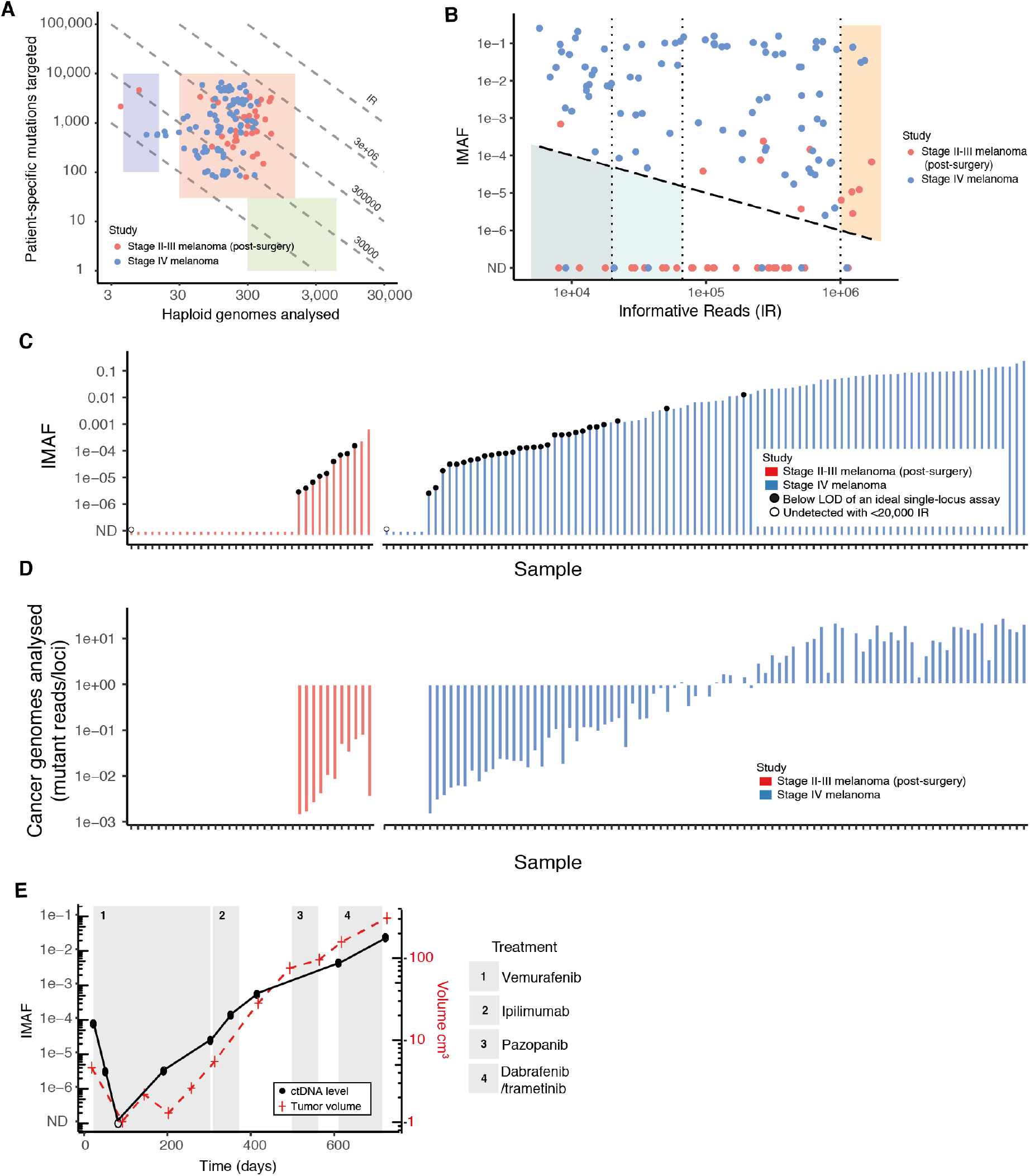
ctDNA detection by INVAR in early and advanced disease. (**A**) Number of haploid genomes analyzed (hGA; calculated as the average depth of unique reads) and the number of mutations targeted, in 144 plasma samples from 66 cancer patients across three cohorts. These were sequenced using custom hybrid-capture panels covering patient-specific mutation lists (Fig. 2B and C) to achieve a median unique depth after read collapsing (hGA) of 185 (Methods) across a median of 628 mutated loci. Dashed diagonal lines indicate the number of targeted loci hGA that yield the indicated IR. (**B**) Two-dimensional representation of detected ctDNA fractions are plotted against the IR for each sample. ctDNA could be detected if its fractional concentration (IMAF) was higher than 2/IR (falling above the dashed line, which is plotted at 1/IR). In some samples, >10^6^ IR were obtained, and ctDNA was detected down to fractions of few ppm (orange shaded region). In some samples, few IR were obtained resulting in limited sensitivity. In our study we used a threshold of 20,000 IR (left-most dotted line), such that samples with undetected ctDNA with fewer than 20,000 IR were called “unclassified” and excluded from the analysis (total of 6 of 144 samples; dark blue shaded region). Samples outside this region had detected ctDNA, or had estimated ctDNA levels below 0.01% (undetected with >20,000 IR; confidence ranges for this value vary for each sample depending on IR). An alternative threshold could be used, for example 66,666 IR, resulting in detection level of 0.003% or 30 ppm (indicated by the second dotted line and the light blue shaded region), increasing the overall detection rates in the cohorts. (**C**) ctDNA fractional levels (IMAF) detected in the samples in this study, shown in ascending order for the two cohorts. Filled circles indicate samples where the number of haploid genomes analyzed would fall below the 95% limit of detection for a perfect single-locus assay given the measured IMAF (Supplementary Methods). Empty circles indicate unclassified samples, i.e. samples for whom ctDNA was not detected (ND) with IR < 20,000. (**D**) The number of copies of the cancer genome detected for each of the samples in the same order as above in part (C), calculated as the number of mutant fragments divided by the number of loci queried (Table S2). (**E**) ctDNA IMAF and tumor volume are plotted over time for one patient with metastatic melanoma over the course of several treatment lines (indicated by shaded boxes). ctDNA was detected to 2.5 ppm during treatment with anti-*BRAF* targeted therapy, when disease volume was approximately 1.3 cm^3^.

In Stage IV melanoma patient samples, ctDNA IMAF values showed a correlation of 0.8 with tumor size assessed by CT imaging (Pearson’s r, P = 6.7 × 10^−10^, Fig. S7B, Table S5), comparable to other studies *(9*, *21)*. Similarly, ctDNA IMAF had a correlation of 0.53 with serum lactate dehydrogenase (LDH), a routinely used clinical marker for monitoring of melanoma (Pearson’s r, P = 2.8 × 10^−4^, Fig. S7C). INVAR analysis was used to monitor ctDNA dynamics in response to treatment, in which the majority of patients received anti-BRAF targeted therapy first line, which resulted in a rapid decline in ctDNA in those patients (Fig. S7D). In one patient (#59) treated with a series of targeted therapies and immunotherapy, ctDNA was detected down to an IMAF of 2.5 ppm, corresponding to a radiological tumor volume of 1.3 cm^3^ (Fig. 6E). Following progression on vemurafenib, patient #59 progressed on multiple other anti-BRAF targeted therapies (pazopanib, dabrafenib and trametinib) and immunotherapy (ipilimumab), corresponding to a constant rise in ctDNA over two years of monitoring (Fig. 6E)

### ctDNA detection post-surgery

To test INVAR in the residual disease setting, we applied INVAR to post-operative samples from 38 patients with resected Stage II-III melanoma recruited in the UK AVAST-M trial. Patient samples were collected up to 6 months after surgery with curative intent. The clinical details of this cohort are given in Fig. S8A. We interrogated a median of 3.6 × 10^5^ IR (IQR 0.64 × 10^5^ to 4.03 × 10^5^) and detected ctDNA to a minimum IMAF of 2.85 ppm, indicated in Fig. 6C. The specificity of this analysis was >0.98 (Fig. S6). In total, ctDNA was detected in samples from 11 of 38 patients (28.9%). ctDNA was detected at higher rates when higher numbers of informative reads were obtained, with ctDNA detected in 10 of 28 (35.7%) cases with >66,666 informative reads (sensitivity of detection of ~30 ppm), 9 of 18 (50%) cases with >250,000 informative reads (detection limit below 10 ppm) and in 5 of 6 (83%) cases with >10^6^ informative reads (Fig. 6B). Samples with no ctDNA detected and few informative reads may indicate limited resolution and would benefit from additional information (more informative reads, obtainable from deeper sequencing or more mutations). A similar approach was previously described in the relative haplotype dosage method by Lo et al. *(22)*. In our case, excluding 3 samples where ctDNA was undetected and had fewer than 20,000 informative reads (detection resolution of 0.01% not reached), ctDNA was detected in 8 of 20 (40%) patients who later recurred and was associated with a strong trend for shorter disease-free interval (6.3 months vs. median not reached with 5 years’ follow-up; Hazard ratio (HR) = 2.08; 95% CI 0.85-5.13, P = 0.11) and overall survival (2.6 years vs. median not reached, P = 0.08). In comparison, a previous analysis of ctDNA detection at 12 weeks after surgery in 161 patients with resected BRAF or NRAS mutant melanoma detected ctDNA in 16.8% of patients who later relapsed *(5)*.

### Sensitive ctDNA monitoring using WES and sWGS

Patient-specific capture panels allow highly sensitive detection of ctDNA, but require prior design of patient specific capture panels. Therefore, we assessed whether INVAR could be applied to standardized workflows such as WES or WGS. This allows the panel design step to be omitted and requires only the patient-specific mutation list from tumor sequencing, which may be performed in parallel with plasma sequencing to save time (Fig. 7A).

**Fig. 7.**
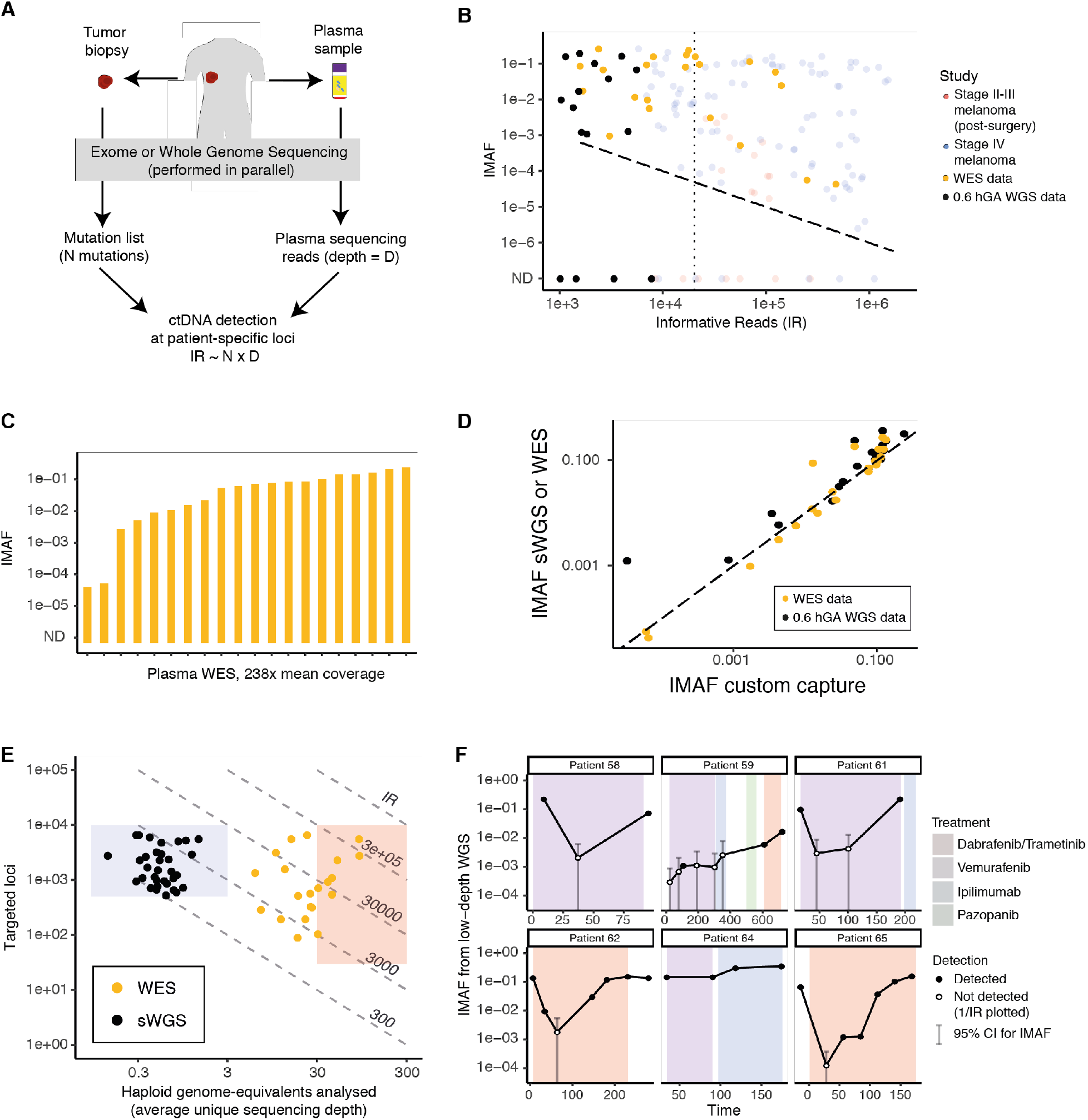
Sensitive detection of ctDNA from WES/WGS data using INVAR. (**A**) schematic overview of a generalized INVAR approach. Tumor (and buffy coat), and plasma samples are sequenced in parallel using whole exome or genome sequencing, and INVAR can be applied to the plasma WES/WGS data using mutation lists inferred from the tumor (and buffy coat) sequencing. (**B**) INVAR was applied to WES data from 21 plasma samples with an average sequencing depth of 238x (before read collapsing), and to WGS data from 33 plasma samples with an average sequencing depth of 0.6x (prior to read collapsing). IMAF values are plotted vs. the number of unique IR for every sample. WES at this depth yielded lower IR compared to the custom capture panel, yet in some cases IR exceeded 10^5^. WGS at low depth yielded <10,000 IR, because mutation lists only spanned the exome based on the extent of tumor sequencing for these cases. The dotted vertical line indicates the 20,000 IR threshold, and the dashed diagonal line indicates 1/IR. (**C**) IMAF observed for the 21 samples analyzed with WES ordered from low to high. ND, not detected. (**D**) IMAFs obtained from plasma WES (gold) and sWGS (black) were compared to the IMAF obtained from the custom capture approach of matched samples, showing correlations of 0.97 and 0.93 (Pearson’s r, P = 1.5 × 10^−13^ and P = 9 × 10^−10^). (**E**) Number of hGA (indicating depth of unique coverage after read collapsing) and mutations targeted by plasma WES and sWGS. Compared to the custom capture approach, both the WES and sWGS samples had fewer hGA and occupy a space further to the left in the two-dimensional space, indicating that INVAR can detect ctDNA from limited data and few genome copies sequenced in a library. (**F**) Longitudinal monitoring of ctDNA levels in plasma of six patients with stage IV melanoma using sWGS data with an average depth of 0.6x, analyzed using INVAR with patient-specific mutation lists (including >500 mutations for each patient, based on WES tumor profiling). Filled circles indicate detection at a specificity level of >0.99 by ROC analysis of the INVAR likelihood (Fig. S9). For other samples, the 95% confidence intervals of the ctDNA level are shown, based on the number of informative reads for each sample (empty circles and bars). ND, not detected.

To test the generalizability of INVAR, we selected samples with IMAF values quantified as being between 4.5 × 10^−5^ and 0.16 using custom-capture sequencing and utilized commercially available exome capture kits to sequence plasma DNA to a median raw depth of 238x. Despite the modest depth of sequencing, we obtained between 1,565 and 473,300 IR using WES (Fig. 7B). We detected ctDNA in all 20 samples tested down to IMAFs as low as 4.34 × 10^−5^ (Fig. 7C), demonstrating that ctDNA can be sensitively detected by INVAR from WES data using patient-specific mutation lists. These IMAF values showed a correlation of 0.97 with custom capture data from the same samples (Pearson’s r, P = 1.5 × 10^−13^, Fig. 7D). Therefore, INVAR is not only highly sensitive when applied to custom capture panels that redundantly sequence up to 10^2^-10^3^ haploid genomes, but also when applied to WES data with a de-duplicated coverage between 10-100x (Fig. 7E).

We hypothesized that ctDNA could be detected and quantified with INVAR from even smaller amounts of input data. Therefore, we performed WGS on libraries from cfDNA of longitudinal plasma samples from a subset of six patients with Stage IV melanoma, to a mean depth of 0.6x (indicated in black in Fig. 7B). For each of those patients we identified >500 patient-specific mutations using WES from each patient’s tumor and buffy coat DNA, generating between 226 and 7,696 IR per sample (median 861, IQR 471-1,559; Fig. 7B) with a “minimum family size” requirement of 1 (i.e. duplicate removal). Despite not leveraging unique molecular barcodes, error rates per trinucleotide were still sufficiently low, with many trinucleotide contexts showing error rates below 1 × 10^−5^ (Fig. S9). Using INVAR on sWGS data, IMAF values as low as 1.1 × 10^−3^ were quantified (Fig. 7F). Compared to custom capture data from the same samples we observed a correlation of 0.93 (Pearson’s r, P = 9 × 10^−10^, Fig. 7D). In samples where ctDNA was not detected, it was possible to estimate the maximum likely IMAF of that sample from the known number of informative reads for each sample, which is indicated by the grey bars in Fig. 7F. Using less than 1x coverage, INVAR can boost the sensitivity utilizing patient-specific mutation lists by up to an order of magnitude compared to copy-number analyses *(15*, *23)*.

These analyses suggest that with a sufficiently large number of tumor -specific mutations, INVAR may yield high sensitivity for ctDNA detection from untargeted sequencing data that can be limited in depth and thus input material obtainable, for example, from dried blood spots *(24)*.

### Extrapolation to higher IR and sensitivity

The sensitivity of INVAR depends on the number of patient-specific mutations identified, and so its effectiveness may be limited in samples with fewer identified mutations, or in cancers with lower mutation rates. Fig. 8A shows the distribution of IR for all the samples in this study, highlighting those with limited sensitivity (<20,000 IR) and those with sensitivity to ppm. Samples with limited sensitivity could be re-analyzed with larger amounts of DNA input/more sequencing, or by designing larger capture panels by identifying additional patient specific mutations through broader-scale sequencing such as WGS of the tumor and buffy coat DNA from that patient. Analyses involving greater IR would render the current background error rates limiting and would therefore require greater error-suppression, such as duplex molecular barcodes *(25)*. However, when increasing the sensitivity of ctDNA beyond the ppm range, sequencing output may become the limiting factor.

**Fig. 8.**
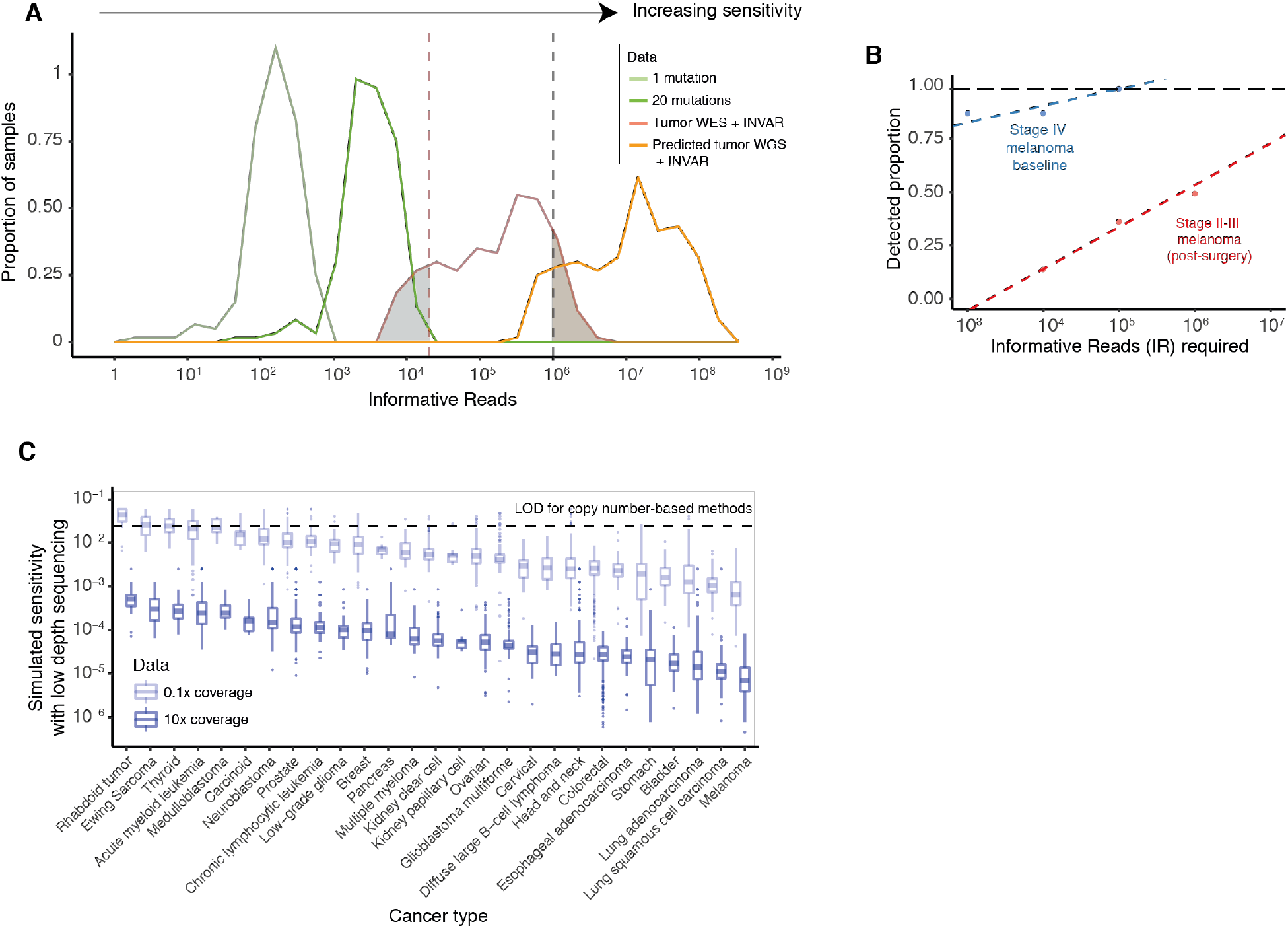
Current limitations and future applications of INVAR. (**A**) The number of informative reads that would be obtainable with different numbers of mutations analyzed, across the cases in these three cohorts. Increasing sensitivity is directly correlated to IR, with the minimal detected ctDNA fraction being 2/IR in the current implementation of INVAR (Methods). The red line shows the distribution of IR obtained with the custom panels covering all mutations identified by tumor WES. Light/dark green lines show the IR generated if 1 or 20 mutations were analyzed for each sample (calculated based on the mean IR per locus). IR could be increased further by using whole genome sequencing (WGS) to guide the design of custom panels (orange curve, extrapolated based on our observed mutation rates in WES). Using mutation lists from WES, samples exceeding 10^6^ IR are shaded in orange, and samples with fewer than 2×10^4^ IR are shaded in blue. (**B**) Detection rates of ctDNA for different numbers of IR sequenced were estimated. There was a linear relationship between IR and detection (R^2^ = 0.95) in the baseline samples of the stage IV melanoma cohort (blue). In stage II-III melanoma post-surgery (red), a linear relationship was observed between IR and detection rate, and the predicted rates of detection of ctDNA was extrapolated. ND, not detected. (**C**) Predicted sensitivities for sWGS plasma analysis of patients with different cancer types, using an average of 0.1x or 10x coverage (equivalent to 0.1 and 10 hGA) and the known mutation rates per Mbp of the genome for different cancer types *(27)*. The limit of detection for ctDNA based on copy number alterations is shown at 3% *(23)*.

We studied the ctDNA levels in the samples analyzed with custom capture in order to estimate the levels of sensitivity that would be required for different clinical applications. We used IMAF values from the clinical samples and plotted the detection rates while varying the numbers of IR and the levels of sensitivity (Fig. 8B). In Stage IV melanoma patients at pre-treatment baseline time points, ctDNA was detected in 100% of cases using 10^5^ IR, whereas up to two orders of magnitude greater sensitivity may be needed to detect ctDNA at high rates following treatment initiation (Fig. S8B). For the population we studied of Stage II-III melanoma patients who underwent surgery, we suggest that even analysis of 10^7^ IR might not be sufficient to detect all patients who ultimately relapse.

## Discussion

In this study, we developed a method for sensitive patient-specific monitoring of ctDNA that leverages the properties of patient-specific sequencing data. This approach mitigates sampling error through aggregation of mutant signal, which is first weighted based on the features of each read and mutation locus, and uses features of cfDNA aside from specific sequence alterations, such as fragment sizes and tumor allele fractions of each mutation. By aggregating signal across 10^2^-10^4^ mutated loci it is possible to detect <0.01 copies of a cancer genome, even when this represents few parts per million of the cfDNA in plasma, 1-2 orders of magnitude lower than previous studies *(6*, *10)*.

We show that INVAR can be applied not only to patient-specific capture panel data to quantify ctDNA to parts per million (Fig. 6), but also to exome sequencing and sWGS data (Fig. 7). Although these latter methods generated fewer informative reads, INVAR detected ctDNA to 50 ppm using WES, and to 0.1% mutant allele fraction using sWGS, over an order of magnitude more sensitive compared to previous methods based on copy-number analysis of sWGS *(23*, *26)*. This level of sensitivity can only be achieved by targeting a sufficiently large number of patient-specific mutations, as increasing input mass alone would not be feasible to this extent (Fig. 2A). Therefore, we assessed whether INVAR would still retain sufficient sensitivity when applied to sWGS data from other cancer types with lower mutation rates than melanoma. We estimated the potential sensitivity for INVAR using sWGS on other cancer types based on their known whole-genome mutation rates *(27)* (Fig. 8C). Using 0.1x WGS coverage, INVAR may yield sensitivities of 10^−1^–10^−3^ for these cancer types, with the potential for higher sensitivity with deeper sequencing.

In summary, patient-specific mutation lists provide an opportunity for highly sensitive monitoring from a range of sequencing data types using methods for signal aggregation, weighting and error-suppression. As tumor sequencing becomes increasingly performed in personalized oncology, patient-specific mutation lists may be leveraged for individualized monitoring using INVAR-like tools.

## Materials and Methods

### Patient cohort

Samples were collected from patients enrolled on the MelResist (REC 11/NE/0312) and AVAST-M studies (REC 07/Q1606/15, ISRCTN81261306) *(28*, *29)*. Consent to enter the studies was taken by a research/specialist nurse or clinician who was fully trained regarding the research. MelResist is a translational study of response and resistance mechanisms to systemic therapies of melanoma, including BRAF targeted therapy and immunotherapy, in patients with stage IV melanoma. AVAST-M is a randomized control trial which assessed the efficacy of bevacizumab in patients with stage IIB-III melanoma at risk of relapse following surgery; only patients from the observation arm were selected for this analysis. The Cambridge Cancer Trials Unit-Cancer Theme coordinated both studies, and demographics and clinical outcomes were collected prospectively. Baseline characteristics for all cohorts are summarized in Table S6.

### Sample collection and processing

Fresh frozen tumor biopsies prior to treatment were collected from patients with Stage IV cutaneous melanoma. Formalin-fixed paraffin-embedded (FFPE) tumor tissue was obtained for the AVAST-M trial. For patients on the AVAST-M study, plasma samples were collected within 12 weeks of tumor resection, with a subsequent sample after 3 months, where available. Longitudinal samples were collected during treatment of patients with stage IV melanoma as part of the MelResist study. Peripheral blood samples were collected at each clinic visit in S-Monovette 9mL EDTA tubes. For plasma collection, samples were centrifuged at 1,600 g for 10 minutes within an hour of the blood draw, and then an additional centrifugation of 20,000 g for 10 minutes was carried out. All aliquots were stored at −80°C.

### Tissue and plasma extraction and quantification

FFPE samples were sectioned into up to 8μm sections, and one H&E stained slide was generated, which was outlined for tumor regions by a histopathologist. Marked tumor regions were macrodissected, and DNA extraction was performed using the QIAamp DNA FFPE Tissue Kit using the standard protocol, except with incubation at 56°C overnight and 500 rpm agitation on a heat block. DNA was eluted twice using 20 μL ATE buffer each time with centrifugation at full speed. Following extraction, DNA repair was performed using the NEBNext® FFPE DNA Repair Mix as per the manufacturer’s protocol. Fresh frozen tissue biopsies were first homogenized prior to DNA extraction, which was performed as follows: up to 30 mg of each fresh frozen tissue biopsy sample was combined with 600 μL RLT buffer, then placed in a Precellys CD14 tube (Bertin Technologies) and homogenized at 6,500 rpm for two bursts of 20 seconds separated by 5 seconds. Subsequently, the Qiagen AllPrep extraction kit as per the manufacturer’s protocol.

Genomic DNA was extracted from up to 1 mL whole blood or buffy coat using the Gentra Puregene Blood Kit (Qiagen) as per the manufacturer’s protocol. Samples were eluted in two rounds of 70 μL buffer AE and incubated for 3 minutes before centrifugation. Up to 4mL of plasma was extracted using the QIAsymphony (Qiagen) with a QIAamp protocol. DNA was eluted in 90 μL elution buffer and stored at −80°C. Plasma samples were extracted using the QIAsymphony instrument (Qiagen) using the 2-4mL QIAamp protocol. For each QIAsymphony batch, 24 samples were extracted, which included a positive and negative control.

Following extraction of fresh frozen, FFPE and genomic DNA, eluted DNA concentration was quantified using a Qubit fluorimeter with a dsDNA broad range assay (ThermoFisher Scientific). To quantify cell-free DNA concentration of plasma DNA eluates, digital PCR was carried out using a Biomark HD (Fluidigm) with a 65bp TaqMan assay for the housekeeping gene RPP30 (Sigma Aldrich) *(7)*. 55 PCR cycles were used. The estimated number of RPP30 DNA copies per μL of eluate was used to determine the cell-free DNA concentration in the original sample.

### Tumor library preparation

FFPE tumor tissue DNA samples (up to 150 ng) and buffy coat DNA samples (75 ng) were sheared to a length of 150bp, using the Covaris LE 220 (Covaris, Massachusetts, USA). The standard Covaris protocol for a final fragment length of 150bp and an input volume of 15μl using the 8 microTUBE-15 AFA Beads Strip V2 was used. After the shearing, the fragmentation pattern was verified using a Bioanalyser (Agilent).

Sequencing libraries were prepared using the ThruPLEX DNA-seq kit (Rubicon). 100ng and 50ng sheared tumor and buffy coat DNA, respectively, were used and the protocol was carried out according to the manufacturer’s instructions. The number of amplification cycles was varied during library preparation according to the manufacturer’s recommendations. Library concentration was determined using qPCR with the Illumina/ROX low Library Quantification kit (Roche). Library fragment sizes were determined using a Bioanalyser (Agilent). After library preparation, exome capture was performed with the TruSeq Exome Library Kit (Illumina), using a 45Mbp exome baitset. Three libraries were multiplexed in one capture reaction and 250ng of each library was used as input. For compatibility with ThruPLEX libraries, the protocol was altered by adding 1μl of i5 and i7 TruSeq HT xGen universal blocking oligos (IDT) during each hybridization step. To compensate for the increased hybridization volume, the volume of CT3 buffer was adjusted to 51 μl. Two rounds of hybridizations were carried out, each lasting for 24 hours. Library QC was performed using qPCR and Bioanalyser, as above. Samples were multiplexed and sequenced with a HiSeq 4000 (Illumina).

Fresh frozen tumor biopsies and matched buffy coat library preparation was performed as described by Varela et al. *(30)* using the SureSelectXT Human All Exon 50 Mb (Agilent) bait set. Samples were multiplexed and sequenced with a HiSeq 2000 (Illumina).

### Tumor mutation calling

For fresh frozen tumor biopsies, mutation calling was performed as described by Varela et al. *(30)*. For FFPE tumor biopsies, mutation calling was performed with Mutect2 with the default settings: --cosmic v77/cosmic.vcf and --dbsnp v147/dbsnp.vcf. To maximize the number of mutations retained, variants achieving Mutect2 pass OR tumor LOD > 5.3 were retained. Mutation calls were filtered as follows:

1. Buffy coat mutant allele fraction equals zero
2. Mutation not in homologous region
3. Mutation not at a multiallelic locus
4. 1000 Genomes ALL and EUR frequency equals zero
5. A minimum unique tumor depth of 5.

In addition, for FFPE data in the melanoma cohort, the filter for C/A errors proposed by Costello et al. *(31)* was applied to suppress C/A artefacts. As a result, we generated patient-specific mutation lists for 47 patients with stage II-IV melanoma. A median of 625 (IQR 411 - 1076) patient-specific mutations were identified per patient (Fig. S2, Table S1). These mutation lists were used both to design custom capture sequencing panels, and as input for the INVAR method.

### Plasma library preparation

Cell-free DNA samples were vacuum concentrated at 30°C using a SpeedVac (ThemoFisher) prior to library preparation where required. The median input into the library was 1652 haploid genomes (IQR 900 – 3013). Whole genome library preparation for plasma cell-free DNA was performed using the Rubicon ThruPLEX Tag-Seq kit. The number of PCR amplification cycles during the ThruPLEX protocol was varied between 7-15 cycles, as recommended by the manufacturer. Following amplification and sample barcoding, libraries were purified using AMPure XP beads (Beckman Coulter) at a 1:1 ratio. Library concentration was determined using the Illumina/ROX low Library Quantification kit (Roche). Library fragment sizes were determined using a Bioanalyser (Agilent).

For the stage IV melanoma cohort, library preparation and sequencing were run in duplicate to assess the technical reproducibility of the experimental and computational method, showing a correlation between IMAF values generated by the INVAR pipeline of 0.97 (Pearson’s r, p-value < 2.2 × 10^−16^). For the early-stage cohorts, input cell-free DNA material was not split and was instead prepared and sequenced as a single sample per time point.

### Custom hybrid-capture panel design and plasma sequencing

Following mutation calling, custom hybrid-capture sequencing panels were designed using Agilent SureDesign software. Between 5 and 9 patients were grouped together per panel in this implementation. Baits were designed with 4-5x density and balanced boosting. 95.5% of the variants had baits successfully designed; bait design was not reattempted for loci that had failed. Custom panels ranged in size between 1.26-2.14 Mb with 120 bp RNA baits. For each panel, mutation classes and tumor allele fractions are shown in Fig. S2 and Table S1.

Libraries were captured either in single or 3-plex (to a total of 1000 ng capture input) using the Agilent SureSelectXT protocol, with the addition of i5 and i7 blocking oligos (IDT) as recommended by the manufacturer for compatibility with ThruPLEX libraries *(32)*. Custom Agilent SureSelectXT baits were used, with 13 cycles of post-capture amplification. Post-capture libraries were purified with AMPure XP beads at a 1:1.8 ratio, then were quantified and library fragment size was determined using a Bioanalyser (Agilent).

### Exome capture sequencing of plasma

For exome sequencing of plasma, the Illumina TruSeq Exome capture protocol was followed. Libraries generated using the Rubicon ThruPLEX protocol (as above) were pooled in 3-plex, with 250ng input for each library. Libraries underwent two rounds of hybridization and capture in accordance with the protocol, with the addition of i5 and i7 blocking oligos (IDT) as recommended by the manufacturer for compatibility with ThruPLEX libraries. Following target enrichment, products were amplified with 8 rounds of PCR and purified using AMPure XP beads prior to QC.

### Plasma sequencing data processing

Cutadapt v1.9.1 was used to remove known 5’ and 3’ adaptor sequences specified in a separate FASTA of adaptor sequences. Trimmed FASTQ files were aligned to the UCSC hg19 genome using BWA-mem v0.7.13 with a seed length of 19. Error-suppression was carried out on ThruPLEX Tag-seq library BAM files using CONNOR *(33)*. The consensus frequency threshold -f was set as 0.9 (90%), and the minimum family size threshold -s was varied between 2 and 5 for characterization of error rates. For custom capture and exome sequencing data, a minimum family size of 2 was used. For sWGS analysis, a minimum family size of 1 was used, i.e. not using molecular barcodes except for where duplicates are present.

To leverage signal across multiple time points, error-suppressed BAM files could be combined using ‘samtools view -ubS - | samtools sort -’ prior to further data processing. In the early-stage melanoma cohort (AVAST-M), where multiple samples were available for the same patient before 6 months post-surgery, sequencing data for each of the samples were merged.

### Low-depth whole-genome sequencing of plasma

For sWGS, 30 libraries were sequenced per lane of HiSeq 4000, achieving a median of 0.6x deduplicated coverage per sample. For these libraries, since the number of informative reads (IR) would limit sensitivity before background errors would become limiting, we used error-suppression with family size 1 for this particular setting. Error rates per trinucleotide were compared between WGS and custom hybrid-capture sequencing data for family size 1, showing a Pearson r of 0.91. WGS data underwent data processing (Supplementary Methods) except the minimum depth at a locus was set to 1, and patient-specific outlier-suppression (Supplementary Methods) was not used because loci with signal vs. loci without signal would only give allele fractions of 0 or 1 given a depth of 0.6x.

### INVAR pipeline

The INVAR pipeline takes BAM files (+/− error-suppression with molecular barcodes), a BED file of patient-specific loci, and a CSV file indicating the tumor allele fraction of each mutation and which patient it belongs to. The pipeline is shown in Fig. S2 and full details are given in the Supplementary Materials. See ‘Data and materials availability’ for code access.

### Imaging

CT imaging was acquired as part of the standard of care from each patient of the stage IV melanoma cohort and was examined retrospectively. Slice thickness was 5 mm in all cases. All lesions with a diameter greater than 5 mm were outlined slice by slice on CT images by an experienced operator, under the guidance of a radiologist, using custom software written in MATLAB (Mathworks, Natick, MA). The outlines were subsequently imported into the LIFEx software *(34)* in NifTI format for processing. Tumor volume was then reported by LIFEx as an output parameter from its texture-based processing module (Table S5).

## Data and materials availability

Raw sequencing data will be made available at the European Genome-phenome archive, accession number EGAS00001002959.

## Supplementary Materials

Materials and Methods

Fig. S1. Flowchart of analysis steps in the INVAR pipeline.

Fig. S2. Tumor mutation list characterization for INVAR.

Fig. S3. Characterization of background error rates.

Fig. S4. Utilizing tumor allelic fraction information and plasma DNA fragment length to enhance ctDNA signal.

Fig. S5. Overview of the INVAR pipeline.

Fig. S6. ROC curves and specificity for all cohorts and data types.

Fig. S7. Characterization of ctDNA levels in advanced melanoma.

Fig. S8. Characterization of IMAF values in the early-stage melanoma cohort.

Fig. S9. Application of INVAR to whole exome sequencing data.

Table S1 Patient-specific mutation lists.

Table S2 Sample library preparation input, QC, and INVAR likelihood ratios – test samples.

Table S3. Sample library preparation input, QC, and INVAR likelihood ratios – control samples.

Table S4 INVAR score thresholds.

Table S5. Tumor volumes for stage IV melanoma cohort

Table S6 Patient baseline characteristics.

## Acknowledgments

The authors would like to thank Catherine Thorbinson, Alex Azevedo, Neera Maroo, from the MelResist and AVAST-M study groups, and the Cambridge Cancer Trial Centre, Addenbrookes Hospital.

## Author contributions

J.C.M.W., K.H. and N.R. wrote the manuscript. J.C.M.W., K.H., S.M., A.R-V., P-Y.C., G.R.B., C.A. and C.G.S. generated data. J.C.M.W., K.H., E.F., J.M., F.Mo., D.C. and N.R. developed the INVAR pipeline. J.C.M.W., K.H., J.M., E.F., A.M., W.Q., F.Ma., J.M. and D.C. analyzed data and performed statistical analysis. A.B.G. and F.A.G. performed imaging analysis. E.B., G.Y., I.H., W.N.C. and D.G. coordinated studies and participated in design. C.P., D.G., A.D., U.M., P.G.C. and N.R. led the MelResist study. P.C.G. and M.M. led the AVAST-M study. C.G.S., C.M., P.G.C., N.R. supervised the project. All authors reviewed and approved the manuscript.

## References and Notes

1. C. Bettegowda, M. Sausen, R. J. Leary, I. Kinde, Y. Wang, N. Agrawal, B. R. Bartlett, H. Wang, B. Luber, R. M. Alani, E. S. Antonarakis, N. S. Azad, A. Bardelli, H. Brem, J. L. Cameron, C. C. Lee, L. a. Fecher, G. L. Gallia, P. Gibbs, D. Le, R. L. Giuntoli, M. Goggins, M. D. Hogarty, M. Holdhoff, S.-M. Hong, Y. Jiao, H. H. Juhl, J. J. Kim, G. Siravegna, D. A. Laheru, C. Lauricella, M. Lim, E. J. Lipson, S. Marie, G. J. Netto, K. S. Oliner, A. Olivi, L. Olsson, G. J. Riggins, A. Sartore-Bianchi, K. Schmidt, L.-M. Shih, S. M. Oba-Shinjo, S. Siena, D. Theodorescu, J. Tie, T. T. Harkins, S. Veronese, T.-L. Wang, J. D. Weingart, C. L. Wolfgang, L. D. Wood, D. Xing, R. H. Hruban, J. Wu, P. J. Allen, C. M. Schmidt, M. a. Choti, V. E. Velculescu, K. W. Kinzler, B. Vogelstein, N. Papadopoulos, L. A. Diaz, Detection of circulating tumor DNA in early- and late-stage human malignancies., Sci. Transl. Med. 6, 224ra24 (2014).

2. J. C. M. Wan, C. Massie, J. Garcia-Corbacho, F. Mouliere, J. D. Brenton, C. Caldas, S. Pacey, R. Baird, N. Rosenfeld, Liquid biopsies come of age: towards implementation of circulating tumour DNA, Nat Rev Cancer 17, 223–238 (2017).

3. I. Garcia-Murillas, G. Schiavon, B. Weigelt, C. Ng, S. Hrebien, R. J. Cutts, M. Cheang, P. Osin, A. Nerurkar, I. Kozarewa, J. A. Garrido, M. Dowsett, J. S. Reis-Filho, I. E. Smith, N. C. Turner, Mutation tracking in circulating tumor DNA predicts relapse in early breast cancer, Sci. Transl. Med. 7(2015), doi:10.1126/scitranslmed.aab0021.

4. J. Tie, Y. Wang, C. Tomasetti, L. Li, S. Springer, I. Kinde, N. Silliman, M. Tacey, H.-L. Wong, M. Christie, S. Kosmider, I. Skinner, R. Wong, M. Steel, B. Tran, J. Desai, I. Jones, A. Haydon, T. Hayes, T. J. Price, R. L. Strausberg, L. A. Diaz Jr, N. Papadopoulos, K. W. Kinzler, B. Vogelstein, P. Gibbs, Circulating tumor DNA analysis detects minimal residual disease and predicts recurrence in patients with stage II colon cancer, Sci. Transl. Med. 8, 346ra92 (2016).

5. R. J. Lee, G. Gremel, A. Marshall, K. A. Myers, N. Fisher, J. A. Dunn, N. Dhomen, P. G. Corrie, M. R. Middleton, P. Lorigan, R. Marais, D. C. Marinescu, Circulating tumor DNA predicts survival in patients with resected high-risk stage II/III melanoma. Ann. Oncol. 29, 490–496 (2018).

6. J. D. Cohen, L. Li, Y. Wang, C. Thoburn, B. Afsari, L. Danilova, C. Douville, A. A. Javed, F. Wong, A. Mattox, R. H. Hruban, C. L. Wolfgang, M. G. Goggins, M. Dal Molin, T.-L. Wang, R. Roden, A. P. Klein, J. Ptak, L. Dobbyn, J. Schaefer, N. Silliman, M. Popoli, J. T. Vogelstein, J. D. Browne, R. E. Schoen, R. E. Brand, J. Tie, P. Gibbs, H.-L. Wong, A. S. Mansfield, J. Jen, S. M. Hanash, M. Falconi, P. J. Allen, S. Zhou, C. Bettegowda, L. A. Diaz, C. Tomasetti, K. W. Kinzler, B. Vogelstein, A. M. Lennon, N. Papadopoulos, Detection and localization of surgically resectable cancers with a multi-analyte blood test, Science (80-.). (2018) (available at http://science.sciencemag.org/content/early/2018/02/15/science.aar3247.abstract).

7. T. Forshew, M. Murtaza, C. Parkinson, D. Gale, D. W. Y. Tsui, F. Kaper, S.-J. S.-J. Dawson, A. M. Piskorz, M. Jimenez-Linan, D. R. Bentley, J. Hadfield, A. P. May, C. Caldas, J. D. Brenton, N. Rosenfeld, Noninvasive Identification and Monitoring of Cancer Mutations by Targeted Deep Sequencing of Plasma DNA, Sci. Transl. Med. 4, 136ra68–136ra68 (2012).

8. M. Murtaza, S.-J. Dawson, K. Pogrebniak, O. M. Rueda, E. Provenzano, J. Grant, S.-F. Chin, D. W. Y. Tsui, F. Marass, D. Gale, H. R. Ali, P. Shah, T. Contente-Cuomo, H. Farahani, K. Shumansky, Z. Kingsbury, S. Humphray, D. Bentley, S. P. Shah, M. Wallis, N. Rosenfeld, C. Caldas, Multifocal clonal evolution characterized using circulating tumour DNA in a case of metastatic breast cancer., Nat. Commun. 6, 8760 (2015).

9. C. Abbosh, N. J. Birkbak, G. A. Wilson, M. Jamal-Hanjani, T. Constantin, R. Salari, J. Le Quesne, D. A. Moore, S. Veeriah, R. Rosenthal, T. Marafioti, E. Kirkizlar, T. B. K. Watkins, N. McGranahan, S. Ward, L. Martinson, J. Riley, F. Fraioli, M. Al Bakir, E. Grönroos, F. Zambrana, R. Endozo, W. L. Bi, F. M. Fennessy, N. Sponer, D. Johnson, J. Laycock, S. Shafi, J. Czyzewska-Khan, A. Rowan, T. Chambers, N. Matthews, S. Turajlic, C. Hiley, S. M. Lee, M. D. Forster, T. Ahmad, M. Falzon, E. Borg, D. Lawrence, M. Hayward, S. Kolvekar, N. Panagiotopoulos, S. M. Janes, R. Thakrar, A. Ahmed, F. Blackhall, Y. Summers, D. Hafez, A. Naik, A. Ganguly, S. Kareht, R. Shah, L. Joseph, A. M. Quinn, P. A. Crosbie, B. Naidu, G. Middleton, G. Langman, S. Trotter, M. Nicolson, H. Remmen, K. Kerr, M. Chetty, L. Gomersall, D. A. Fennell, A. Nakas, S. Rathinam, G. Anand, S. Khan, P. Russell, V. Ezhil, B. Ismail, M. Irvin-Sellers, V. Prakash, J. F. Lester, M. Kornaszewska, R. Attanoos, H. Adams, H. Davies, D. Oukrif, A. U. Akarca, J. A. Hartley, H. L. Lowe, S. Lock, N. Iles, H. Bell, Y. Ngai, G. Elgar, Z. Szallasi, R. F. Schwarz, J. Herrero, A. Stewart, S. A. Quezada, K. S. Peggs, P. Van Loo, C. Dive, C. J. Lin, M. Rabinowitz, H. J. W. L. Aerts, A. Hackshaw, J. A. Shaw, B. G. Zimmermann, the TRACERx consortium, the PEACE consortium, C. Swanton, Phylogenetic ctDNA analysis depicts early stage lung cancer evolution, Nature 22364, 1–25 (2017).

10. A. M. Newman, A. F. Lovejoy, D. M. Klass, D. M. Kurtz, J. J. Chabon, F. Scherer, H. Stehr, C. L. Liu, S. V Bratman, C. Say, L. Zhou, J. N. Carter, R. B. West, G. W. Sledge Jr., J. B. Shrager, B. W. Loo Jr., J. W. Neal, H. A. Wakelee, M. Diehn, A. A. Alizadeh, Integrated digital error suppression for improved detection of circulating tumor DNA, Nat Biotechnol 34, 547–55 (2016).

11. B. R. McDonald, T. Contente-Cuomo, S.-J. Sammut, A. Odenheimer-Bergman, B. Ernst, N. Perdigones, S.-F. Chin, M. Farooq, R. Mejia, P. A. Cronin, K. S. Anderson, H. E. Kosiorek, D. W. Northfelt, A. E. McCullough, B. K. Patel, J. N. Weitzel, T. P. Slavin, C. Caldas, B. A. Pockaj, M. Murtaza, Personalized circulating tumor DNA analysis to detect residual disease after neoadjuvant therapy in breast cancer, Sci. Transl. Med. 11, eaax7392 (2019).

12. C. Turnbull, Introducing whole-genome sequencing into routine cancer care: the Genomics England 100 000 Genomes Project, Ann. Oncol. 29, 783–784 (2018).

13. J. Phallen, M. Sausen, V. Adleff, A. Leal, C. Hruban, J. White, V. Anagnostou, J. Fiksel, S. Cristiano, E. Papp, S. Speir, T. Reinert, M.-B.W. Orntoft, B. D. Woodward, D. Murphy, S. Parpart-Li, D. Riley, M. Nesselbush, N. Sengamalay, A. Georgiadis, Q. K. Li, M. R. Madsen, F. V. Mortensen, J. Huiskens, C. Punt, N. van Grieken, R. Fijneman, G. Meijer, H. Husain, R. B. Scharpf, L. A. Diaz, S. Jones, S. Angiuoli, T. Ørntoft, H. J. Nielsen, C. L. Andersen, V. E. Velculescu, Direct detection of early-stage cancers using circulating tumor DNA, Sci. Transl. Med. 9(2017) (available at http://stm.sciencemag.org/content/9/403/eaan2415.abstract).

14. K. C. A. Chan, J. K. S. Woo, A. King, B. C. Y. Zee, W. K. J. Lam, S. L. Chan, S. W. I. Chu, C. Mak, I. O. L. Tse, S. Y. M. . Leung, G. Chan, E. P. Hui, B. B. Y. Ma, R. W. K. Chiu, S.-F. Leung, A. C. van Hasselt, A. T. C. Chan, Y. M. D. Lo, Analysis of Plasma Epstein–Barr Virus DNA to Screen for Nasopharyngeal Cancer, N. Engl. J. Med. 377, 513–522 (2017).

15. F. Mouliere, D. Chandrananda, A. M. Piskorz, E. K. Moore, J. Morris, L. B. Ahlborn, R. Mair, T. Goranova, F. Marass, K. Heider, J. C. M. Wan, A. Supernat, I. Hudecova, I. Gounaris, S. Ros, M. Jimenez-linan, J. Garcia-corbacho, K. Patel, O. Østrup, S. Murphy, M. D. Eldridge, D. Gale, G. D. Stewart, J. Burge, W. N. Cooper, M. S. Van Der Heijden, C. E. Massie, C. Watts, P. Corrie, S. Pacey, K. M. Brindle, R. D. Baird, M. Mau-sørensen, Enhanced detection of circulating tumor DNA by fragment size analysis, Sci. Transl. Med. 4921, 1–14 (2018).

16. I. Kinde, J. Wu, N. Papadopoulos, K. W. Kinzler, B. Vogelstein, Detection and quantification of rare mutations with massively parallel sequencing, Proc. Natl. Acad. Sci. U. S. A. 108, 9530–5 (2011).

17. M. Jamal-Hanjani, G. A. Wilson, S. Horswell, R. Mitter, O. Sakarya, T. Constantin, R. Salari, E. Kirkizlar, S. Sigurjonsson, R. Pelham, S. Kareht, B. Zimmermann, C. Swanton, Detection of ubiquitous and heterogeneous mutations in cell-free DNA from patients with early-stage non-small-cell lung cancer, Ann. Oncol. 27, 862–867 (2016).

18. H. R. Underhill, J. O. Kitzman, S. Hellwig, N. C. Welker, R. Daza, D. N. Baker, K. M. Gligorich, R. C. Rostomily, M. P. Bronner, J. Shendure, Fragment Length of Circulating Tumor DNA, PLoS Genet. 12, 426–37 (2016).

19. F. Mouliere, B. Robert, E. Arnau Peyrotte, M. Del Rio, M. Ychou, F. Molina, C. Gongora, A. R. Thierry, T. Lee, Ed. High Fragmentation Characterizes Tumour-Derived Circulating DNA, PLoS One 6, e23418 (2011).

20. H. C. Fan, Y. J. Blumenfeld, U. Chitkara, L. Hudgins, S. R. Quake, Analysis of the size distributions of fetal and maternal cell-free DNA by paired-end sequencing, Clin. Chem. 56, 1279–1286 (2010).

21. A. M. Newman, S. V Bratman, J. To, J. F. Wynne, N. C. W. Eclov, L. a Modlin, C. L. Liu, J. W. Neal, H. a Wakelee, R. E. Merritt, J. B. Shrager, B. W. Loo, A. a Alizadeh, M. Diehn, An ultrasensitive method for quantitating circulating tumor DNA with broad patient coverage, Nat. Med. 20, 548–54 (2014).

22. Y. M. D. Lo, K. C. A. Chan, H. Sun, E. Z. Chen, P. Jiang, F. M. F. Lun, Y. W. Zheng, T. Y. Leung, T. K. Lau, C. R. Cantor, R. W. K. Chiu, Maternal Plasma DNA Sequencing Reveals the Genome-Wide Genetic and Mutational Profile of the Fetus, Sci. Transl. Med. 2(2010) (available at http://stm.sciencemag.org/content/2/61/61ra91.full).

23. V. A. Adalsteinsson, G. Ha, S. S. Freeman, A. D. Choudhury, D. G. Stover, H. A. Parsons, G. Gydush, S. C. Reed, D. Rotem, J. Rhoades, D. Loginov, D. Livitz, D. Rosebrock, I. Leshchiner, J. Kim, C. Stewart, M. Rosenberg, J. M. Francis, C.-Z. Zhang, O. Cohen, C. Oh, H. Ding, P. Polak, M. Lloyd, S. Mahmud, K. Helvie, M. S. Merrill, R. A. Santiago, E. P. O’Connor, S. H. Jeong, R. Leeson, R. M. Barry, J. F. Kramkowski, Z. Zhang, L. Polacek, J. G. Lohr, M. Schleicher, E. Lipscomb, A. Saltzman, N. M. Oliver, L. Marini, A. G. Waks, L. C. Harshman, S. M. Tolaney, E. M. Van Allen, E. P. Winer, N. U. Lin, M. Nakabayashi, M.-E. Taplin, C. M. Johannessen, L. A. Garraway, T. R. Golub, J. S. Boehm, N. Wagle, G. Getz, J. C. Love, M. Meyerson, Scalable whole-exome sequencing of cell-free DNA reveals high concordance with metastatic tumors, Nat. Commun. 8, 1324 (2017).

24. K. Heider, J. C. M. Wan, J. Hall, S. Boyle, I. Hudecova, D. Gale, W. N. Cooper, P. G. Corrie, J. D. Brenton, C. G. Smith, N. Rosenfeld, Detection of ctDNA from dried blood spots after DNA size selection, BioRxiv (2019).

25. M. W. Schmitt, S. R. Kennedy, J. J. Salk, E. J. Fox, J. B. Hiatt, L. A. Loeb, Detection of ultra-rare mutations by next-generation sequencing, Proc. Natl. Acad. Sci. 109, 14508–14513 (2007).

26. J. Belic, M. Koch, P. Ulz, M. Auer, T. Gerhalter, S. Mohan, K. Fischereder, E. Petru, T. Bauernhofer, J. B. Geigl, M. R. Speicher, E. Heitzer, Rapid Identification of Plasma DNA Samples with Increased ctDNA Levels by a Modified FAST-SeqS Approach, Clin. Chem. 61, 838–849 (2015).

27. M. S. Lawrence, P. Stojanov, P. Polak, G. V. Kryukov, K. Cibulskis, A. Sivachenko, S. L. Carter, C. Stewart, C. H. Mermel, S. A. Roberts, A. Kiezun, P. S. Hammerman, A. McKenna, Y. Drier, L. Zou, A. H. Ramos, T. J. Pugh, N. Stransky, E. Helman, J. Kim, C. Sougnez, L. Ambrogio, E. Nickerson, E. Shefler, M. L. Cortés, D. Auclair, G. Saksena, D. Voet, M. Noble, D. DiCara, P. Lin, L. Lichtenstein, D. I. Heiman, T. Fennell, M. Imielinski, B. Hernandez, E. Hodis, S. Baca, A. M. Dulak, J. Lohr, D.-A. Landau, C. J. Wu, J. Melendez-Zajgla, A. Hidalgo-Miranda, A. Koren, S. A. McCarroll, J. Mora, R. S. Lee, B. Crompton, R. Onofrio, M. Parkin, W. Winckler, K. Ardlie, S. B. Gabriel, C. W. M. Roberts, J. A. Biegel, K. Stegmaier, A. J. Bass, A. Garraway, M. Meyerson, T. R. Golub, D. A. Gordenin, S. Sunyaev, E. S. Lander, G. Getz, Mutational heterogeneity in cancer and the search for new cancer-associated genes, Nature 499, 214–218 (2013).

28. P. G. Corrie, A. Marshall, J. A. Dunn, M. R. Middleton, P. D. Nathan, M. Gore, N. Davidson, S. Nicholson, C. G. Kelly, M. Marples, S. J. Danson, E. Marshall, S. J. Houston, R. E. Board, A. M. Waterston, J. P. Nobes, M. Harries, S. Kumar, G. Young, P. Lorigan, Adjuvant bevacizumab in patients with melanoma at high risk of recurrence (AVAST-M): Preplanned interim results from a multicentre, open-label, randomised controlled phase 3 study, Lancet Oncol. 15, 620–630 (2014).

29. P. G. Corrie, A. Marshall, P. D. Nathan, P. Lorigan, M. Gore, S. Tahir, G. Faust, C. G. Kelly, M. Marples, S. J. Danson, E. Marshall, S. J. Houston, R. E. Board, A. M. Waterston, J. P. Nobes, M. Harries, S. Kumar, A. Goodman, A. Dalgleish, A. Martin-Clavijo, S. Westwell, R. Casasola, D. Chao, A. Maraveyas, P. M. Patel, C. H. Ottensmeier, D. Farrugia, A. Humphreys, B. Eccles, G. Young, E. O. Barker, C. Harman, M. Weiss, K. A. Myers, A. Chhabra, S. H. Rodwell, J. A. Dunn, M. R. Middleton, Adjuvant bevacizumab for melanoma patients at high risk of recurrence: Survival analysis of the AVAST-M trial, Ann. Oncol. 29, 1843–1852 (2018).

30. I. Varela, P. Tarpey, K. Raine, D. Huang, C. K. Ong, H. Davies, D. Jones, M. Lin, J. Teague, G. Bignell, A. Butler, J. Cho, G. L. Dalgliesh, D. Galappaththige, C. Hardy, M. Jia, C. Latimer, K. W. Lau, J. Marshall, S. Mclaren, A. Menzies, L. Mudie, L. Stebbings, A. David, L. F. a Wessels, S. Richard, R. J. Kahnoski, J. Anema, Exome sequencing identifies frequent mutation of the SWI / SNF complex gene PBRM1 in renal carcinoma, Nature 469, 539–542 (2011).

31. M. Costello, T. J. Pugh, T. J. Fennell, C. Stewart, L. Lichtenstein, J. C. Meldrim, J. L. Fostel, D. C. Friedrich, D. Perrin, D. Dionne, S. Kim, S. B. Gabriel, E. S. Lander, S. Fisher, G. Getz, Discovery and characterization of artifactual mutations in deep coverage targeted capture sequencing data due to oxidative DNA damage during sample preparation, Nucleic Acids Res. 41, 1–12 (2013).

32. Rubicon Genomics, Targeted Capture of ThruPLEX® Libraries with Agilent SureSelect®XT Target Enrichment System (available at rubicongenomics.com/wp-content/uploads/2016/11/RDM-152-002-SureSelectXT.pdf).

33. University of Michigan, Connor - METHODS (2016) (available at https://github.com/umich-brcf-bioinf/Connor/blob/master/doc/METHODS.rst).

34. C. Nioche, F. Orlhac, S. Boughdad, S. Reuze, M. Soussan, C. Robert, C. Barakat, I. Buvat, A freeware for tumor heterogeneity characterization in PET, SPECT, CT, MRI and US to accelerate advances in radiomics, J. Nucl. Med. 58, 1316 (2017).

