## Supplementary Material for "High-sensitivity monitoring of ctDNA by patient-specific sequencing panels and integration of variant reads"

### Contents

**Supplementary Methods**

**Supplementary Figures**

| | |
|---|---|
| Fig. S1 | Flowchart of analysis steps in the INVAR pipeline |
| Fig. S2 | Tumor mutation list characterization for INVAR |
| Fig. S3 | Characterization of background error rates |
| Fig. S4 | Utilizing tumor allelic fraction information and plasma DNA fragment length to enhance ctDNA signal |
| Fig. S5 | Overview of the INVAR pipeline |
| Fig. S6 | ROC curves and specificity for all cohorts and data types |
| Fig. S7 | Characterization of ctDNA levels in advanced melanoma |
| Fig. S8 | Characterization of IMAF values in the early-stage melanoma cohort |
| Fig. S9 | Application of INVAR to whole exome sequencing data |

**Supplementary Table Legends**

| | |
|---|---|
| Table S1 | Patient-specific mutation lists |
| Table S2 | Sample library preparation input, QC, and INVAR likelihood ratios – test samples |
| Table S3 | Sample library preparation input, QC, and INVAR likelihood ratios – control samples |
| Table S4 | INVAR score thresholds |
| Table S5 | Tumor volumes for stage IV melanoma cohort |
| Table S6 | Patient baseline characteristics |

**Supplementary References**

### Supplementary Methods

INVAR data processing

SAMtools mpileup 1.3.1 was used at patient-specific loci based on a BED file of mutations, with the following settings: --ff UNMAP, –q 40 (mapping quality), –Q 20 (base quality), –x, --d 10,000, then multiallelic calls were split using BCFtools 1.3.1. Next, all TSV files were annotated with 1,000 Genomes SNP data, COSMIC data, and trinucleotide context using a custom Python script. Output files were then concatenated, compressed, and read into R. First, based on prior knowledge from tumor sequencing data, all loci were annotated per patient with being either: patient-specific (present in patient's tumor) or non-patient-specific (not present in patient's tumor, or individual does not have cancer). Data points were excluded if MQSB < 0.01 (mapping quality / strand bias). Since each non-patient-specific sample contains the loci from multiple patients, every non-patient-specific sample may control for all other patients analyzed with the same sequencing panel or method (excluding loci that are shared between individuals).

INVAR data filters I

The following filters were applied to INVAR data:

1. Loci that showed mutant signal in >10% of the non-patient-specific (patient-control) samples were blacklisted. For custom capture and exome sequencing data, we required the mean background error rate in each locus to be <1% mutant allele fraction, otherwise the locus was classed as noisy. The proportion of loci that were blacklisted with these filters ranged from 0.21%-3.53% (Fig. S3D). Patient samples may be used to characterize the noise per locus (at loci that did not belong to them), since 99.8% of mutations were private to each patient. Control sample QC data are shown in Table S3.
2. Mutation signal had to be represented in both the F and R read of that read pair (Fig. S3C). The resulting error-suppression is analogous to tools that merge paired-end reads*(1)*.

INVAR data annotation

After data filtering, data was annotated with both locus noise filter and trinucleotide error rate. Since the locus noise filter is limited by the number of control samples and cfDNA molecules at that locus, we also assessed trinucleotide error rate. Trinucleotide error rates were determined from the region up to 10bp either side of every patient-specific locus (excluding patient-specific locus itself), and data was pooled by trinucleotide context. After pooling data in this manner, a median of $3.0 \times 10^8$ informative reads (or deduplicated reads) per trinucleotide context were analyzed. Trinucleotide error rate was calculated as a mismatch rate for each specific mutation context. If a trinucleotide context had zero mutant deduplicated reads, the error rate was set to the reciprocal of the number of IR/deduplicated reads in that context.

In addition, each data point was annotated with the cfDNA fragment size of that read using a custom Python script. Then, to eliminate outlier signal that was not consistent with the remainder of that patient's loci, we performed patient-specific outlier suppression (Fig. 4 B and C, Fig. S3G). The data is now error-suppressed (both by read-collapsing and bespoke methods for patient-specific sequencing data) and annotated with parameters required for signal-enrichment (by features of ctDNA sequencing) for the INVAR method.

INVAR data filters II - patient-specific outlier-suppression

Patient-specific sequencing data consists of informative reads at multiple known patient-specific loci, providing the opportunity to compare mutant allele fractions across loci as a means of error-suppression. The distribution of signal across loci potentially allows for the identification of noisy loci not consistent with the overall signal distribution. Each locus was tested for the probability of having observed mutant reads given the average signal across all loci (Fig. 4 B and C, Fig. S3G). A locus observed with significantly greater signal than the remainder of the loci might be attributed to noise at that locus, contamination, or a mis-genotyped SNP locus. The possibility of a mis-genotyped SNP becomes increasingly likely when a large number of mutated loci are targeted by INVAR.

For each sample, the IMAF was determined across all loci passing pre-INVAR data processing filters with mutant allele fraction at that locus of <0.25, similar to that outlined by Phallen et al.*(2)*, who used a similar threshold, and Abbosh et al.*(3)*, who used a threshold of 0.20. Loci with signal >0.25 mutant allele fraction were not included in the calculation because (i) in the residual disease setting, loci would not be expected to have such high mutant allele fractions (unless they were mis-genotyped SNPs), and (ii) if the true IMAF of a sample is >0.25, when a large number of loci are tested, they will show a distribution of allele fractions such that detection is supported by having many low allele fraction loci with signal.

Based on the ctDNA level of the sample, the binomial probability of observing each individual locus given the IMAF of that sample was calculated. Loci with a Bonferroni corrected P-value <0.05 (corrected for the number of loci interrogated) were excluded in that sample, thereby suppressing outliers. As a result of outlier-suppression, background noise was reduced to 33% in control samples, while retaining 96.1% of signal in patient samples (Fig. 4C). By correcting the P-value threshold for the number of loci tested, this filter can be applied to data with a variable number of mutations targeted per patient, enabling analysis of samples from patients with cancer types with both high and low mutation rates.

Statistical detection method for INVAR

We developed a statistical method to model the number of mutant reads at multiple patient-specific loci, incorporating prior information available from patient-specific sequencing, such as the background error of the trinucleotide context, the tumor allele fraction at the locus, and fragment length. This approach aggregates signal across multiple patient-specific mutations following error-suppression. For each locus, we test the significance of the number of mutant reads given the trinucleotide error rate of that context. Trinucleotide error rates were used instead of locus-specific error rates in order to determine a more accurate estimation of background error rates to $10^{-7}$ (Fig. S3E).

Tumor allele fractions and trinucleotide error rates were considered as follows: Denote $AF_i$ as the tumor mutant allele fraction at locus $i$, $e_i$ as the background error in the context of locus $i$, and let $p$ be an estimate of ctDNA content in that sample for the INVAR pipeline. A random read at locus $i$ can be observed to be mutant either if it arose from a mutant molecule, or an incorrectly sequenced wild type DNA molecule. This occurs with probability $q_i$:

$$q_i = AF_i \cdot (1 - e_i) \cdot p + (1 - AF_i) \cdot e_i \cdot p + e_i \cdot (1 - p) \qquad (1)$$

Testing for the presence of ctDNA is now equivalent to testing the statistical hypothesis $H_0: p = 0$. Assuming the number of observed mutant reads is independent between loci, the following likelihood function can be produced:

$$L(p; M, AF, e) = \prod_{i=1}^{n} \prod_{j=1}^{R_i} {q_i}^{M_{ij}} (1 - q_i)^{1 - M_{ij}} \quad (2)$$

where $M_{ij}$ is the indicator for a mutation in read *j* of locus *i*, and $R_i$ is the number of reads in locus *i*. The above method allows weighting of signal by tumor allele fraction, which we confirm influences plasma mutation representation in patient samples with early stage and advanced disease (Fig. 4D), and in the spike-in dilution series from one patient (Fig. S4A).

Each sequencing read provides fragment size information (Fig. S4B), which may be used to separate mutant from wild-type molecules and produce an enrichment in ctDNA (Fig. 4E). Probability weighing was preferred over size selection to avoid allelic loss at ultra-low allele fractions, suggested by Fan et al.*(4)* in the non-invasive prenatal testing setting. Therefore, read length information can also be incorporated into the likelihood. The method for read length distribution of mutant and wild-type fragments estimation is given in the section *Estimation of read length distribution for INVAR*. This approach is in contrast to size-selection and may be considered as a size-weighting step alongside tumor AF weighting that was performed above. Fragment sizes for each sequencing read may be incorporated to the INVAR method. To do so, let $L_{ji}$ be the length of read *j* at locus *i*. The likelihood can be written as:

$$L(p; M, L, AF, e) = \prod_{i=1}^{n} \prod_{j=1}^{R_i} P(m_{ij}, l_{ij} | e, AF, p) \quad (3)$$

Assuming that the read length and mutation status are independent given the source of the read (ctDNA or non-tumor cfDNA), we can factor the likelihood as follows:

$$\begin{aligned} L(p; M, L, AF, e) &= \prod_{i=1}^{n} \prod_{j=1}^{R_i} P(m_{ij}, l_{ij} | z_{ij} = 0) \cdot P(z_{ij} = 0) + P(m_{ij}, l_{ij} | z_{ij} = 1) \cdot P(z_{ij} = 1) \\ &= \prod_{i=1}^{n} \prod_{j=1}^{R_i} P(m_{ij} | z_{ij} = 0) \cdot p^0(l_{ij}) \cdot (1 - p) + P(m_{ij} | z_{ij} = 1) \cdot p^1(l_{ij}) \cdot p \\ &= \prod_{i=1}^{n} \prod_{j=1}^{R_i} {e_i}^{m_{ij}} \cdot (1 - e_i)^{1 - m_{ij}} \cdot p^0(l_{ij}) \cdot (1 - p) + g_i^{m_{ij}} \cdot (1 - g_i)^{1 - m_{ij}} \cdot p^1(l_{ij}) \\ &\cdot p \quad (4) \end{aligned}$$

where $z_{ij}$ is the indicator that read *j* of locus *i* came from ctDNA, $p^k(l_{ij}) = P(l_{ij} | z_{ij} = k)$, and $g_i = AF_i \cdot (1 - e_i) + (1 - AF_i) \cdot e_i$. The above method weights the signal in one sample based on both fragment length of mutant and wild-type reads from all other samples, though in this implementation of INVAR, we set the weight of all wild-type size bins to be equal, thereby neglecting size information from wild-type reads.

Lastly, a score is generated for each sample through aggregation of signal across all patient-specific loci in that sample using the Generalized Likelihood Ratio test (GLRT). The GLRT directly compares the likelihood under the null hypothesis against the likelihood under the maximum likelihood estimate of *p*:

$$\lambda(p_0) = \frac{L(p_0; M, L, AF, e)}{L(\hat{p}; M, L, AF, e)}$$

(5)

The higher the value of the likelihood ratio, the greater the evidence for ctDNA presence in a sample. Classification of samples was performed based on comparison of likelihood ratios between patient and control samples.

Likelihood ratio threshold determination

Other patients were used to control for one another at non-shared loci (Fig. 3D). Only samples run on the same sequencing panel (i.e. same custom sequencing panel design), with the same error-suppression setting and targeting the same mutation list were used to control for one another.

In order to determine an accurate threshold for the likelihood ratio (LR) based on controls, reads from each control sample were iteratively resampled with replacement 10 times, and the GLRT script was run. To minimize the risk of any patient-specific contamination of signal at non-patient-specific control loci (through *de novo* mutations overlapping with patient-specific sites), only samples with patient-specific IMAF <1% were used as controls for determination of the cut-point. Control samples were required to have at least as many IR as the patient sample with the fewest IR in that cohort, otherwise they were excluded.

Based on the LR distribution in patient controls and patient samples, the cut-off for LR was determined for each cohort using the 'OptimalCutpoints' package in R*(5)*, maximizing sensitivity and specificity using the 'MaxSnSp' setting. Based on the LRs per cohort, an analytical specificity was determined for each cohort (Fig. S6, Table S4).

Assessment of specificity in healthy individuals

26 healthy individuals' cfDNA from plasma were analyzed using the stage IV melanoma cohort. These samples were treated as 'patient' samples, and so had no influence on the filters in the pipeline and were not used for the determination of LR thresholds. After determination of the LR thresholds (described above), the LRs from healthy individuals' samples were assessed for false positive detection of ctDNA. For each of the INVAR applications (custom capture, WES and sWGS), the clinical specificity values in healthy individuals were determined (Fig. S6, Table S4).

Estimation of ctDNA content per sample for likelihood ratio determination

In this section we derive an Expectation Maximization (EM) algorithm to estimate $p$ as part of the INVAR method. If we treat the tumor of origin $z_{ij}$ as a latent variable, and assume that it is known, the joint likelihood of Z, M ($m_{ij}$ is the indicator for a mutation in read $j$ of locus $i$), L ($l_{ij}$ is the length of read $i$ of locus $j$), AF ($AF_i$ is the tumor allele fraction at locus $i$), e ($e_i$ is the background error in the context of locus $i$) can be written as:

$$L(p; Z, M, L, AF, e) = \prod_{i=1}^{n} \prod_{j=1}^{R_i} [e_i^{m_{ij}} \cdot (1-e_i)^{1-m_{ij}} \cdot p^0(l_{ij}) \cdot (1-p)]^{1-z_{ij}} \cdot [g_i^{m_{ij}} \cdot (1-g_i)^{1-m_{ij}} \cdot p^1(l_{ij}) \cdot p]^{z_{ij}}$$

Where $g_i = AF_i \cdot (1-e_i) + (1-AF_i) \cdot e_i$. The log-likelihood is linear in $z_{ij}$, so taking the expectation of the likelihood amounts simply to replacing the $z_{ij}$ with their expectation at stage $l$, $z_{ij}^l = E(z_{ij} | m_{ij}, l_{ij}, p_l)$, where $p_l$ is the best estimate of $p$ at iteration $l$. We can thus use EM to find a maximum likelihood estimate for $p$, by iteratively maximizing the likelihood with respect to $p$, and taking the expectation of the likelihood with respect to $z_{ij}$. An estimate for $p_l$ is obtained by taking the derivative with respect to $p_l$ and equating it to zero:

$$p_l = \frac{\sum_{i=1}^{n} \sum_{j=1}^{R_i} z_{ij}^l}{\sum_{i=1}^{n} R_i}$$

The above is simply the expected proportion of reads from ctDNA at stage $l$. Bayes' theorem can be used to compute $z_{ij}^l$:

$$z_{ij}^l = P(z_{ij} = 1 | m_{ij}, l_{ij}, p_l)$$
$$= \frac{P(m_{ij}|z_{ij} = 1) \cdot p^1(l_{ij}) \cdot p}{P(m_{ij}|z_{ij} = 1) \cdot p^1(l_{ij}) \cdot p + P(m_{ij}|z_{ij} = 0) \cdot p^0(l_{ij}) \cdot (1 - p)}.$$

By substituting the respective probabilities, we obtain:

$$z_{ij}^l = \frac{g_i^{m_{ij}}(1 - g_i)^{1-m_{ij}} p^1(l_{ij}) p}{g_i^{m_{ij}}(1 - g_i)^{1-m_{ij}} p^1(l_{ij}) p + e_i^{m_{ij}}(1 - e_i)^{1-m_{ij}} p^0(l_{ij})(1 - p)}$$

The algorithm proceeds by alternating the maximization of $p$, and the expectation of the $z_{ij}$.

Estimation of read length distribution for INVAR

Size-weighting with INVAR depends on first having a known distribution of sizes of mutant and wild-type reads against which to perform weighting. In order to estimate the read length distribution with the greatest accuracy, we used all wild type and mutant reads from all samples in that panel, leaving out the sample being tested (i.e. leave-one-out approach), and we used kernel density estimation to smooth the respective probabilities.

The size distributions from each of the studied cohorts are shown in Fig. S4, and the enrichment ratios for each size range are shown in Fig. 4E. We demonstrated that the early stage melanoma cohort had a significantly different size profile from the advanced stage melanoma cohort, which had a significantly greater proportion of di-nucleosomal fragments despite downsampling of data to a similar number of reads (Fig. S4C). Thus, the fragment length distribution of mutant and wildtype fragments was assessed separately in the two cohorts, and data was smoothed with a Gaussian kernel with a default setting of 0.25 (Fig. S4D).

To estimate the probability that a read is of length $l$, given that the cell of origin is wild type, $P(L = l | z = 0)$, we used all of the wild-type reads from each pooled dataset. For both of the data sets, we used the R function "density", with a Gaussian kernel, to smooth the estimated probabilities, and obtained a density estimate $\hat{f}(l|Z = z)$. Finally, to estimate $P(L = l|z = z)$, we integrated the respective density:

$$P(L = l|Z = z) = \int_{l-0.5}^{l+0.5} \hat{f}(t|Z = z) dt$$

Smoothing the size distribution estimates is important in datasets where data is sparse to avoid assigning too large a weight to any given mutant fragment.

Calculation of informative reads (IR)

The number of informative reads (IR) for a sample is the product of the number of mutations targeted (i.e. length of the mutation list) and the number of haploid genomes analyzed by sequencing (hGA, equivalent to the deduplicated coverage following read-collapsing). Thus, the limit of detection for every sample can be calculated based on 1/IR (with adjustment for sampling mutant molecules based on binomial probabilities). For non-detected samples, the 1/IR value provides an estimate for the upper limit of ctDNA in that sample; this allows quantification of samples even if no mutant molecules are present, and is utilized in Fig. 7F to define the upper confidence limits to ~$10^{-4}$ using sWGS data. Also, samples with limited sensitivity can be identified and classified as a 'low-sensitivity' or 'non-evaluable' group, where the INVAR method is limited by the number of IR (Fig. 8A). In this study, we aimed to quantify ctDNA with sensitivity greater than other methods, and classified samples with non-detected ctDNA with <20,000 IR as low-sensitivity and thus non-evaluable. Across the cohorts in this study, 4 patients were non-evaluable with these criteria.

Calculation of integrated mutant allele fraction (IMAF)

To quantify ctDNA across multiple mutated loci, we calculated an 'integrated mutant allele fraction', as follows:

a) For each trinucleotide context in a sample, the deduplicated depth-weighted mean allele fraction across all patient-specific loci was calculated

b) The background error rate per trinucleotide context in control data was subtracted from the mean allele fraction calculated in (a). Trinucleotide contexts with negative mutant allele fraction after subtraction were set to zero.

c) The mean background-subtracted allele fraction was taken across the trinucleotide contexts, weighted by the deduplicated depth in each trinucleotide context.

Experimental spike-in dilution series

Plasma DNA from one patient with a total of 5,073 patient-specific variants was serially diluted 10-fold each step in a pool of plasma cfDNA from 11 healthy individuals (Seralab) to give a dilution series spanning 1-100,000x. Library preparation was performed, as described in Methods, with 50ng input per dilution. In order to interrogate a sufficiently large number of molecules in the dilution series to assess sensitivity, the lowest dilution (100,000x) was generated in triplicate. The healthy control cfDNA pools were included as control samples for the determination of locus error rate to identify and exclude potential SNP loci (Fig. 5A).

Given the relationship between tumor allele fraction and plasma mutation representation (Fig. S4A), any smaller panel for INVAR should be based on clonal mutations with highest priority, with lower allele fractions included only if plasma sequencing data is sufficiently broad. Thus, we iteratively sampled the data with replacement from each of the dilution series sequencing libraries (with 50 iterations), and then selected the top N mutations (spanning 1 to 5,000 mutations). The locus with the highest mutant allele fraction was the *BRAF* V600E mutation. After downsampling the number of loci, outlier-suppression was repeated on all samples except for the single BRAF *V600E* locus data (Fig. 5B).

### Supplementary Figures

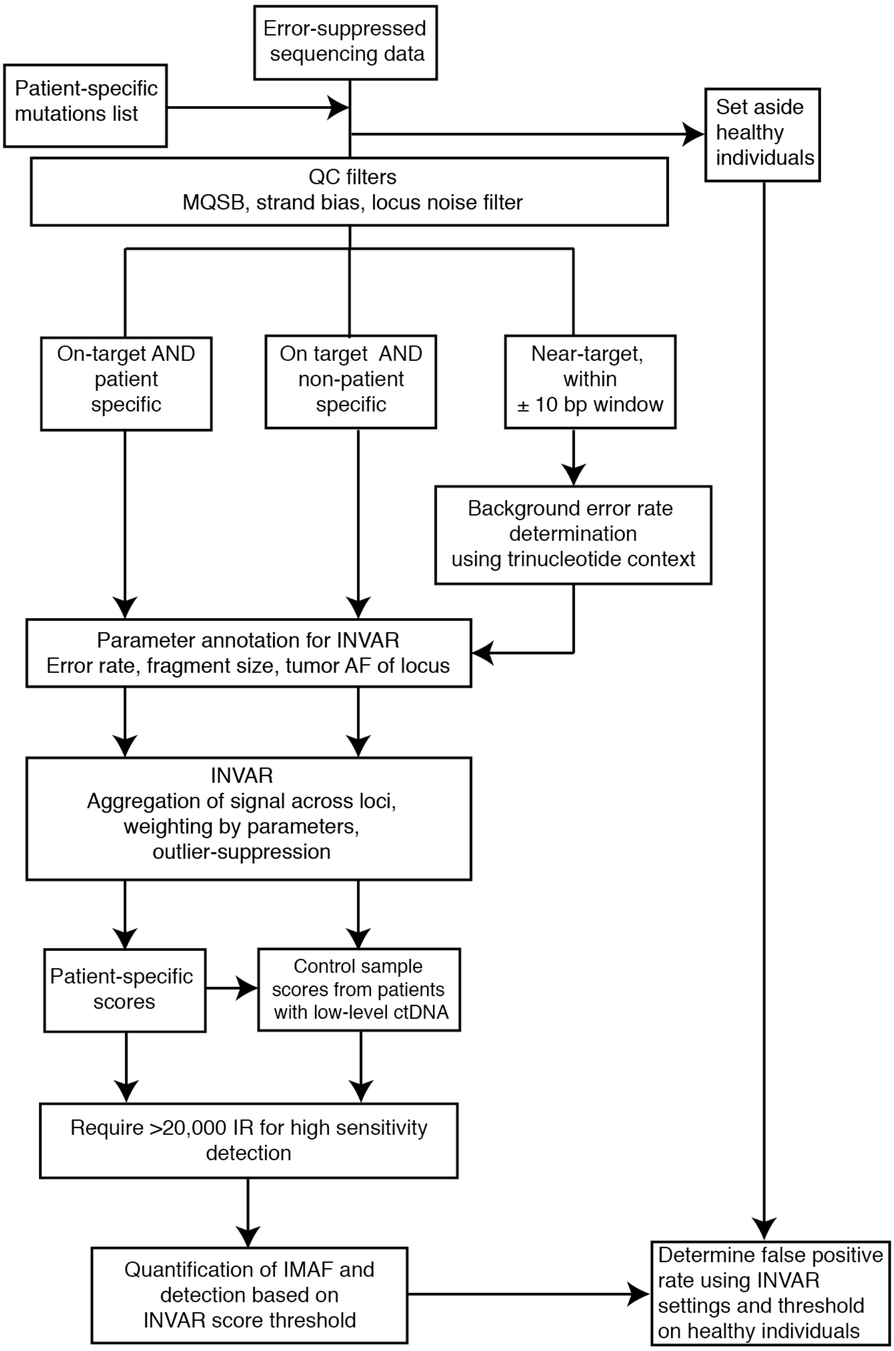


**Fig. S1. Flowchart of analysis steps in the INVAR pipeline.**

Integration of variant reads workflow. INVAR utilizes plasma sequencing data and requires a list of patient-specific mutations, which may be derived from tumor or plasma sequencing. Filters are applied to sequencing data, then the data is split into: patient-specific (locus belonging to that patient), non-patient-specific (locus not belonging to that patient), and near-target (bases within 10 bp of all patient-specific loci). Patient-specific and non-patient-specific data are annotated with features that influence the probability of observing a real mutation. Outlier-suppression is applied to identify mutant signal inconsistent with the overall level of patient-specific signal. Next, signal is aggregated across all loci, considering annotated features, to generate an INVAR score per sample. Based on non-patient-specific samples, an INVAR score threshold is determined using ROC analysis for each cohort. Healthy control samples separately undergo the same steps to establish a specificity value for each cohort.

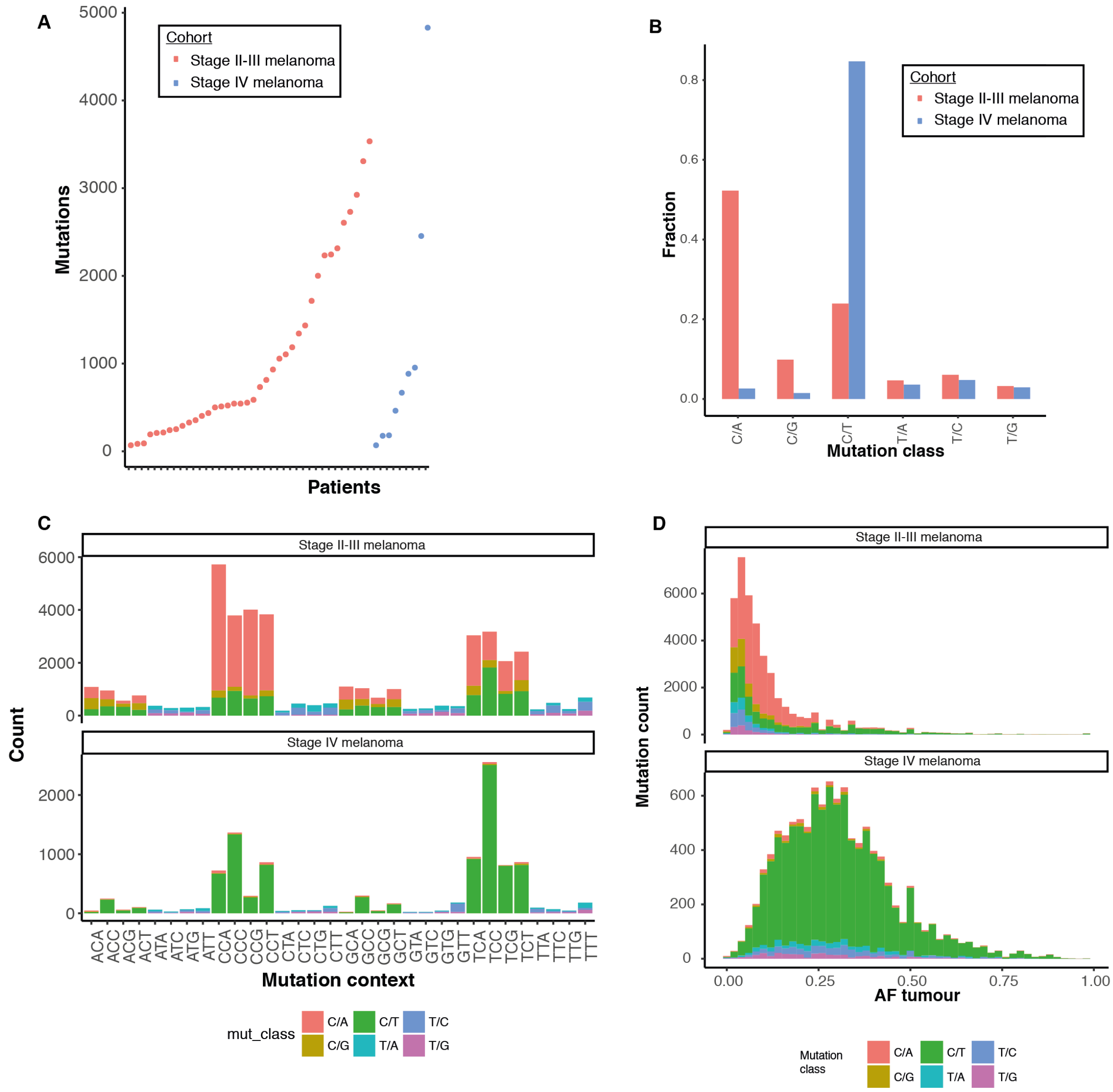


**Fig. S2. Tumor mutation list characterization for INVAR.**

300 (**A**) Number of somatic mutations per patient, ordered by cohort. (**B**) Frequency of each mutation class included in each panel design. (**C**) Mutation counts by trinucleotide context, colored by mutation class. (**D**) Distribution of tumor mutation allele fractions per cohort colored by mutation class.

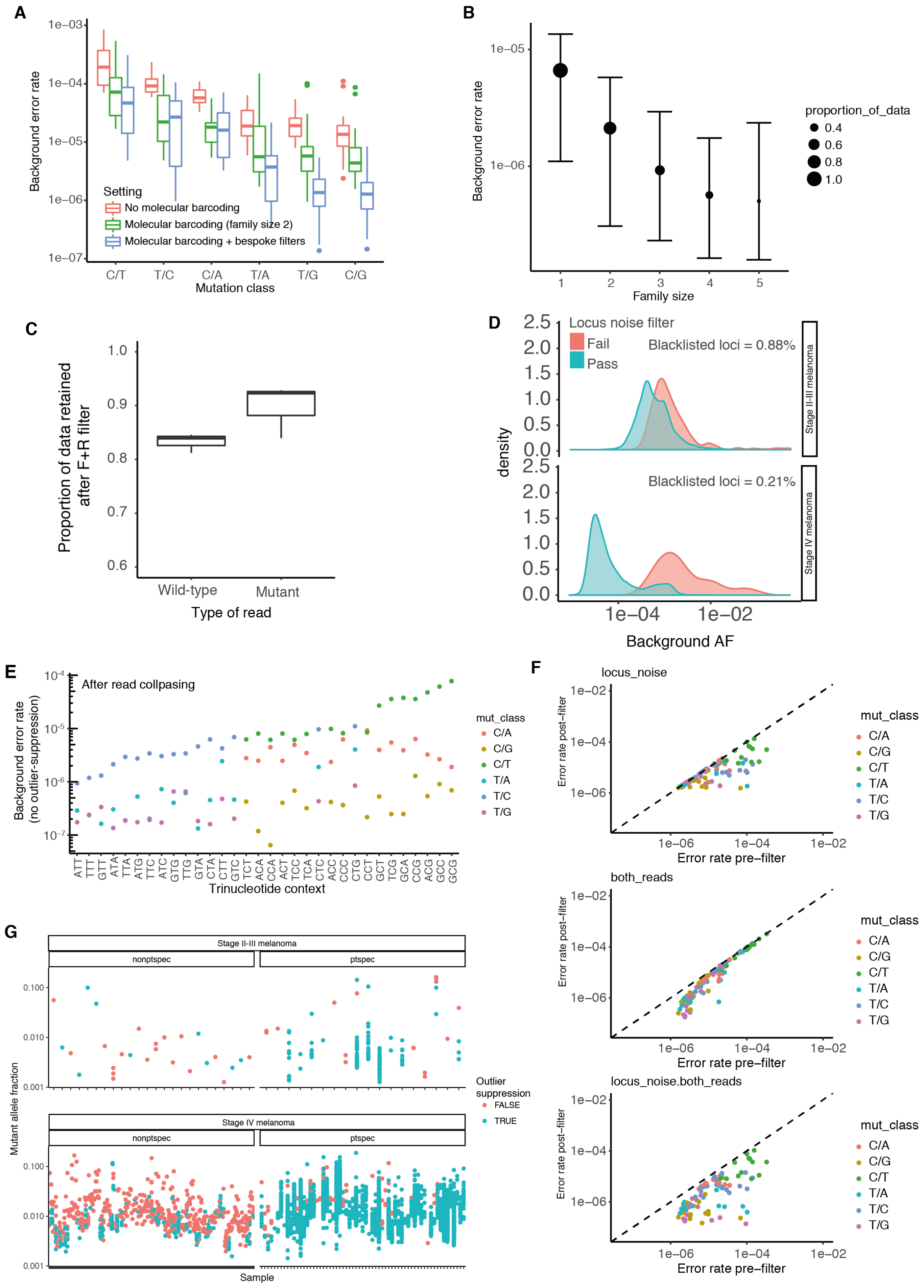


305 **Fig. S3. Characterization of background error rates.**

(**A**) Error suppressed (minimum family size of 2) and non-error suppressed background error rates, with and without bespoke INVAR filters. Background error rates were calculated by aggregating all non-reference bases across all considered bases. To assess background error rate, a 10 bp window either side of the patient-specific loci was used, excluding the patient-specific locus itself ('near-target', Supplementary Methods). (**B**) Overall background error rates resulting from different minimum family size requirements, and the proportion of read families retained with each setting. (**C**) Effect of requiring forward and reverse reads at a locus; a median of 84.0% of the wild-type reads and a median of 92.4% of the mutant reads were retained with this filter. (**D**) Background error rates were characterized per locus based on all reads generated from control samples, split by cohort. The loci that passed the locus noise filter are shown in blue, loci that did not pass the filter are shown in red. The proportions of loci blacklisted by this filter are indicated at the top right. (**E**) Error rates by trinucleotide context and mutation class following data filtering but before applying the patient specific outlier suppression. Error rates can vary by more than an order of magnitude within the same mutation class, highlighting the need to assess loci with respect to their trinucleotide context. This requires the aggregation of large amounts of data across multiple loci. (**F**) For each trinucleotide context, background error rates (per trinucleotide) are plotted before and after each background error filter, highlighting the collective benefit of each of the error filters. (**G**) Outlier suppression filter: Raw data-points for both cohorts (showing patient and control samples), with the outlier-suppressed data points indicated in red (Details in Supplementary Methods).

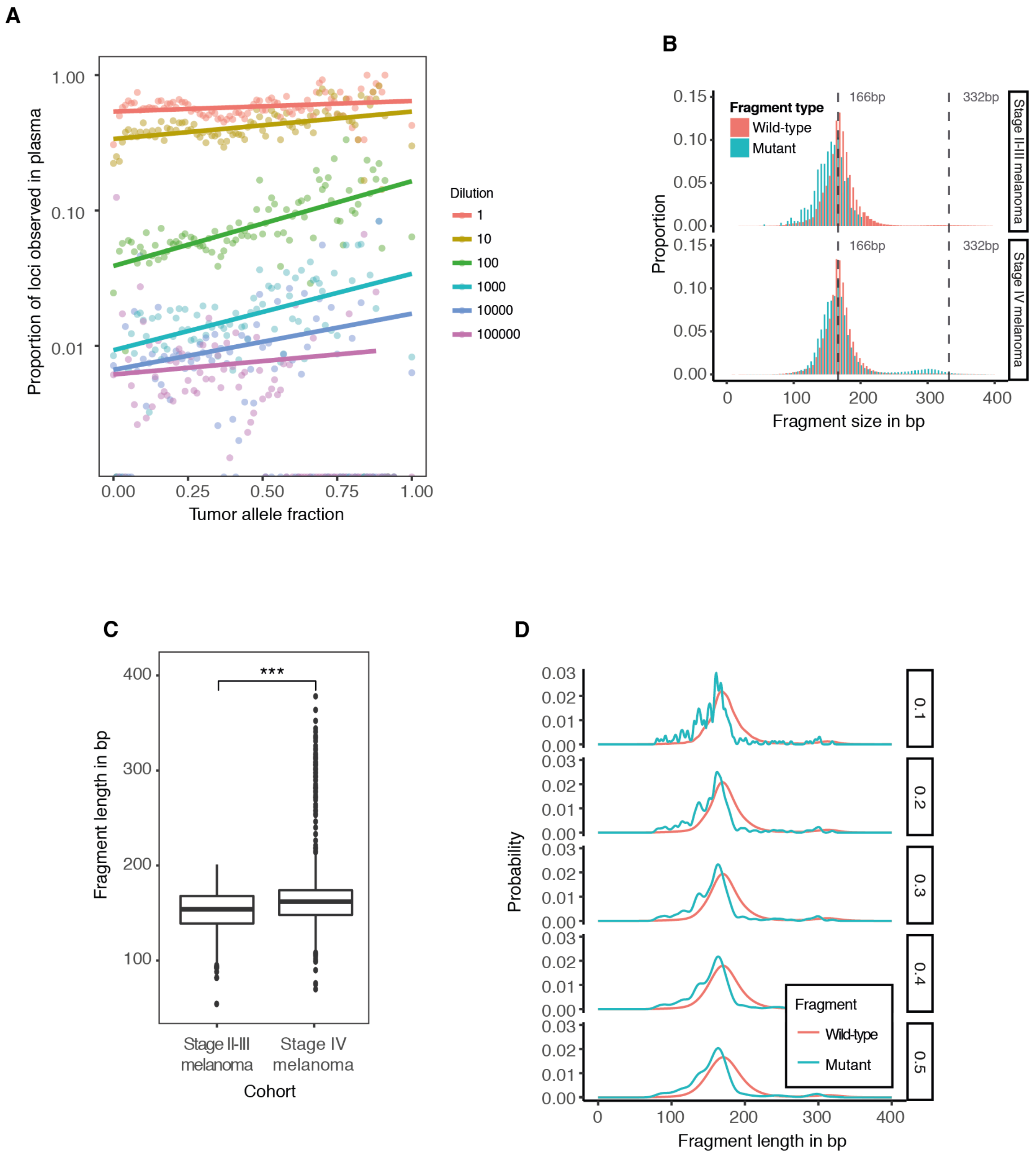


**Fig. S4. Utilizing tumor allelic fraction information and plasma DNA fragment length to enhance ctDNA signal.**

(**A**) Comparison of tumor and plasma mutant allele fractions. Using error-suppressed data, tumor loci were grouped into bins of 0.01 mutant allele fraction, and the proportion of loci observed in plasma was determined for different levels of a dilution series. The dilution level of the spike-in dilution series is indicated by each color. At each dilution level, there is a positive correlation between the tumor allele fraction and proportion of loci observed in plasma. (**B**) For each cohort, size profiles were generated for mutant and wild-type fragments. (**C**) Comparison of mutant fragment distributions between cohorts. These were compared using a two-sided Wilcoxon rank test after downsampling the number of mutant reads to match for both cohorts. (**D**) The distributions of fragment sizes for different levels of smoothening, used to assign weights to fragment sizes (Supplementary Methods).

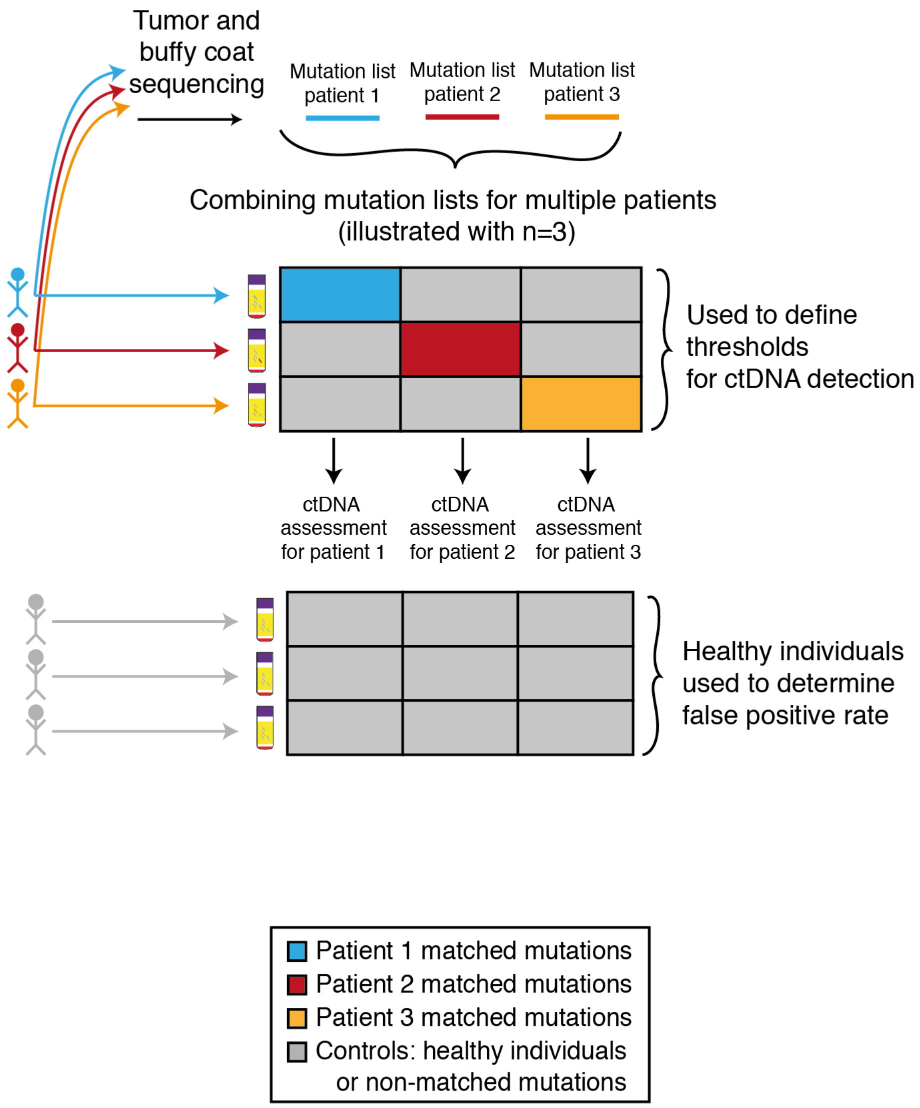


**Fig. S5. Overview of the INVAR pipeline.**

INVAR leverages patients to control for one another and uses separate healthy controls. In this study, individual mutation lists are generated from tumor and buffy coat sequencing. Each locus of interest is sequenced in the matched patient, and in additional patients from the same cohort for whom this locus was not found to be mutated in the tumor or buffy coat analysis. This can be done by applying a generic panel to all samples (such as WES/WGS, Fig. 7), or by combining multiple patient-specific mutation lists into a combined custom panel that is sequenced across multiple patients (Fig. 6). For each patient, INVAR aggregates the sequencing information across the loci of the patient-specific mutation list. Data from other patients in those loci ('non-matched mutations') are used to determine background mutation rates and detection cut-offs (Supplementary Methods). Additional samples from healthy individuals are analyzed by the same panels, this data was used to assess the false positive rates in healthy individuals.

**A**

| Cohort | Analytical spec. | Specificity |
|---|---|---|
| Stage IV melanoma | 95.4% | 96.7% |
| Stage II-III melanoma | 98.4% | N/A |
| WES plasma | 96.1% | 95.8% |
| sWGS plasma | 100% | 97.6% |

**B**

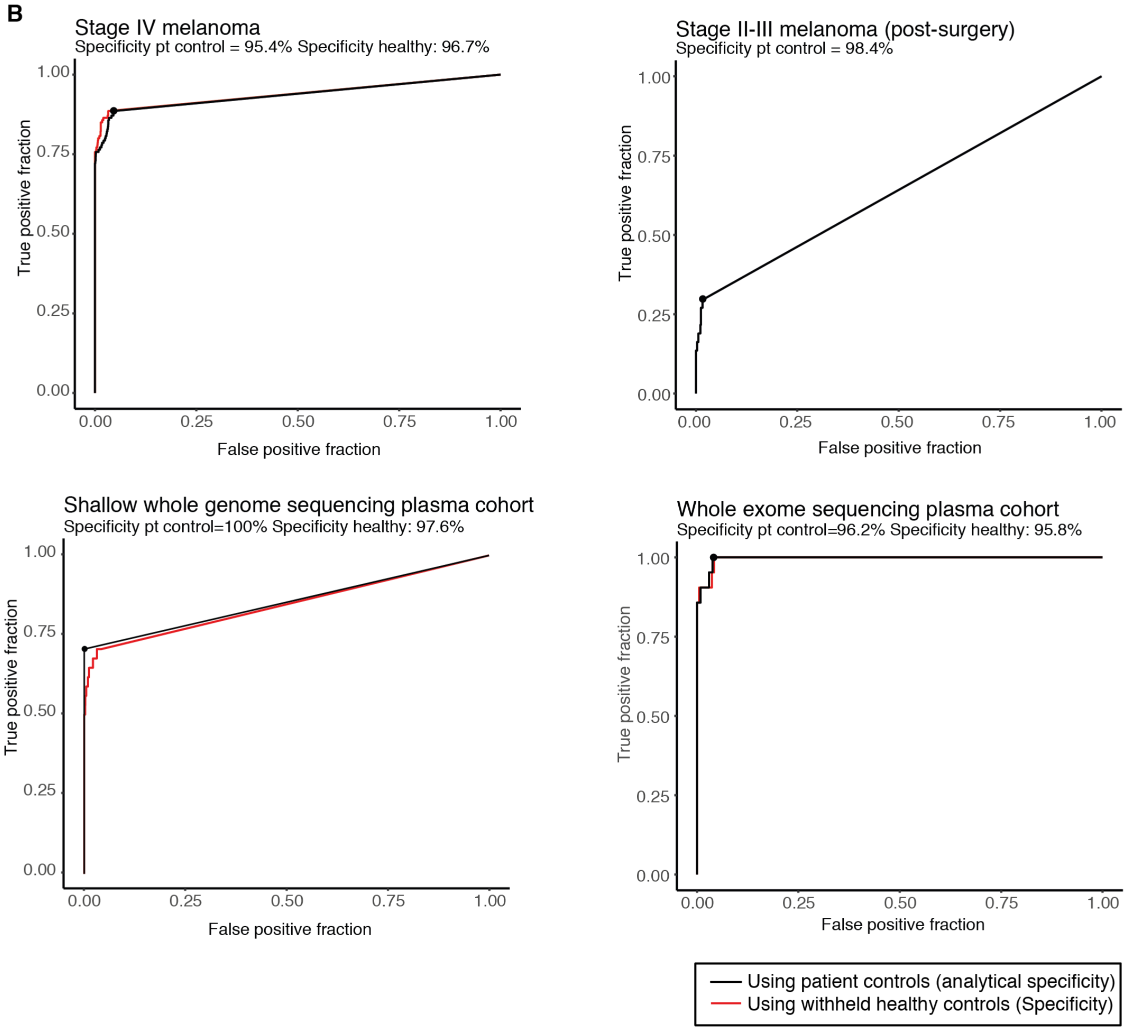


**Fig. S6. ROC curves and specificity for all cohorts and data types**.

(**A**) For all cohorts, the analytical (based on patient control samples) and clinical specificity values (based on healthy individuals) are given. No healthy individuals were tested on the custom capture panel of the stage II-III melanoma cohort. (**B**) For all cohorts, ROC analyses were performed against patient-controls (black) and healthy individuals (red). For the stage II-III melanoma (post-surgery) cohort, our analysis was blinded to outcome, and patients who did not relapse within 5 years were also included in the ROC analysis; thus, the maximal possible sensitivity for this cohort (as defined) was the fraction of relapsing patients (18/33=54.5%). INVAR detected 9 out of 18 patients who relapsed (ROC showing sensitivity at 9/33=27.3%).

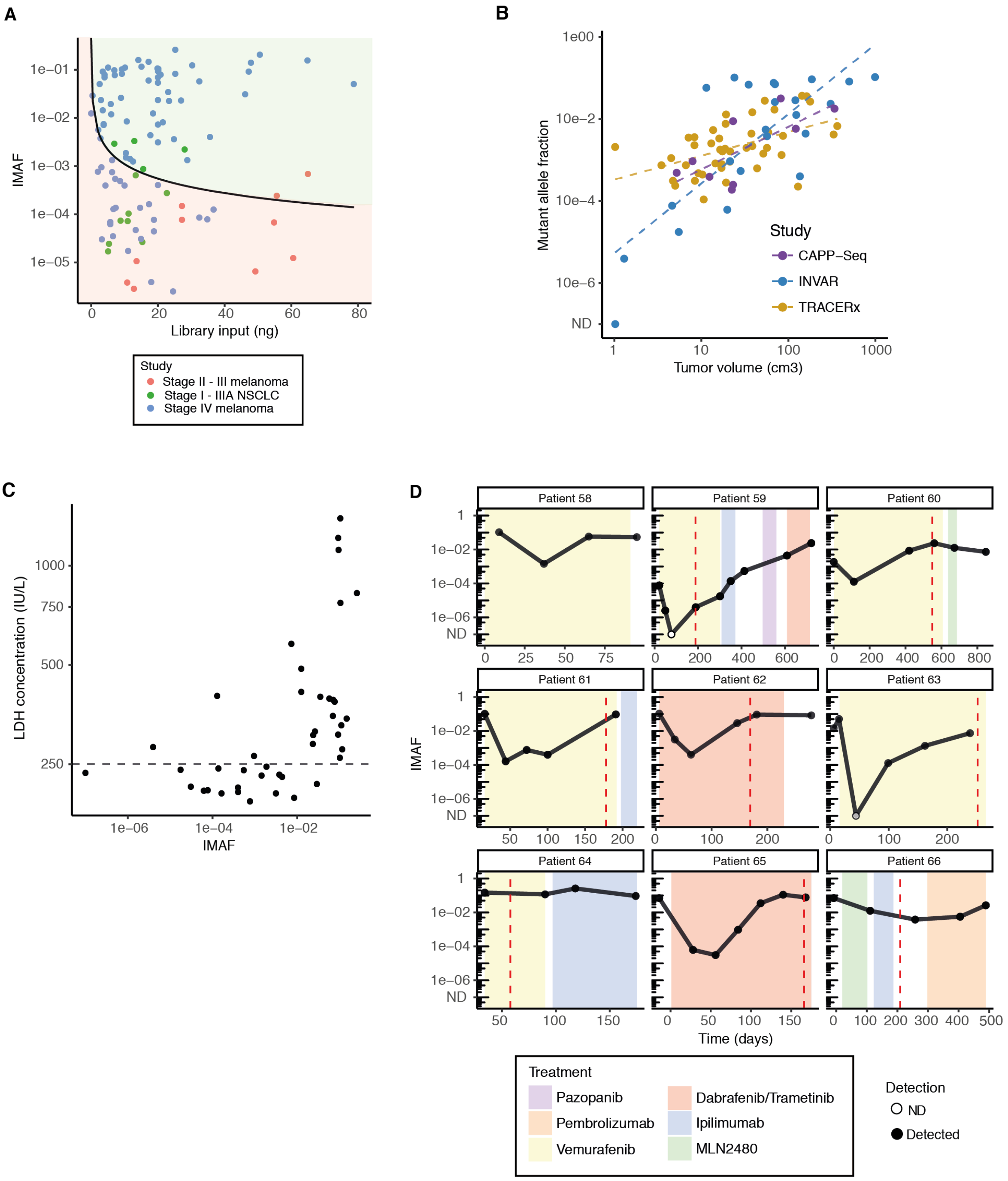


**Fig. S7. Characterization of ctDNA levels in advanced melanoma.**

(**A**) Comparison of input mass and IMAF observed. For each library with detected ctDNA, the DNA input mass for library preparation is plotted against the IMAF determined by INVAR. The black line indicates the threshold below which a perfect single-locus assay would have <95% sensitivity, based on the likelihood that no mutant copies would be sampled given the expected number of mutant copies in the sample. In this study, 48% of samples would not be detectable using a perfect single-locus assay with the plasma DNA input amounts used. (**B**) Comparison between ctDNA and tumor volume in our study (Pearson's r = 0.67, P = 0.0002) and in previous publications measuring multiple mutations per patient in NSCLC, using CAPP-Seq*(6)*, and using multiplexed PCR in the TRACERx cohort*(3)*. The relationship between tumor volume and ctDNA level was steeper in this study than in previous analyses. This may be due to detection of ctDNA at lower concentrations using INVAR, which may have been missed or over-estimated by other assays. (**C**) Relationship between serum lactate dehydrogenase and IMAF in advanced stage melanoma patients. A Pearson correlation score of 0.46 was observed (P = 0.0058). A dashed line is drawn at 250IU/L, the upper limit of normal for LDH. (**D**) Longitudinal ctDNA profiles for advanced melanoma patients. IMAF values are plotted over time per patient, using error-suppressed individualized sequencing data. Vertical dashed lines indicate time of radiological progression.

A

| | Total | |
|---|---|---|
| | N | % |
| **Characteristics** | | |
| | 38 | |
| **Sex** | | |
| Male | 17 | 45 |
| Female | 21 | 55 |
| **Breslow thickness** | | |
| <=2.0mm | 16 | 42 |
| >2-4.0mm | 8 | 21 |
| >4.0mm | 12 | 32 |
| Unknown | 2 | 5 |
| **Ulceration** | | |
| Present | 9 | 24 |
| Absent | 23 | 60 |
| Unknown | 6 | 16 |
| **Disease stage** | | |
| II | 8 | 21 |
| IIIA | 4 | 10 |
| IIIB | 17 | 45 |
| IIIC | 9 | 24 |
| **N classification** | | |
| II (No or N/A) | 8 | 21 |
| III (N1a and N2a) | 6 | 16 |
| III (other N) | 24 | 63 |
| **ECOG performance status** | | |
| 0 | 35 | 92 |
| 1 | 3 | 8 |
| **BRAF and NRAS mutation** | | |
| BRAF mutation | 19 | 50 |
| NRAS mutation | 8 | 21 |
| BRAF and NRAS WT | 10 | 26 |
| BRAF WT and NRAS not tested | 1 | 3 |
| **Trial arm** | | |
| Observation | 38 | 100 |
| **Time from latest surgery to trial entry in weeks** | | |
| Median (Range) | 11 (4-12) | |

B

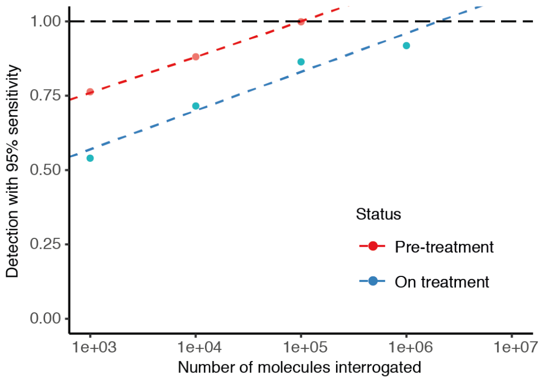


**Fig. S8. Characterization of IMAF values in the early-stage melanoma cohort.**

(**A**) Summary table of patient characteristics for the stage II-III resected melanoma cohort (n = 38). (**B**) We estimated the detection rates of ctDNA for different levels of IR (Supplementary Methods). We observe a linear relationship ($R^2 = 0.95$) between the number of IR and detection rate in the baseline samples of the stage IV melanoma cohort. ctDNA was detected in 100% of baseline samples with $10^5$ IR (red), whereas following the initiation of treatment, $10^6$-$10^7$ IR are needed to detect all longitudinal samples (blue), reflecting the lower levels of ctDNA.

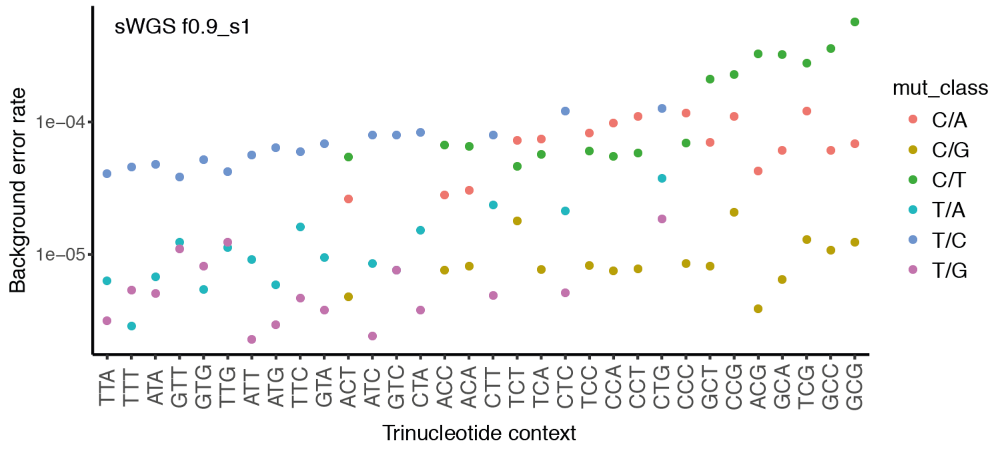


400 **Fig. S9. Application of INVAR to whole genome sequencing data.**

Background error rates per trinucleotide context for sWGS data. Data was analyzed with INVAR using a minimum family size of 1 when performing read collapsing.

### Supplementary Table Legends

Tables are attached separately.

**Table S1** Patient-specific mutation lists. This table contains all patient-specific mutation lists for patients in this study. The following cohorts are represented: AVASTM (stage II-III melanoma) and MELR (stage IV melanoma). Mutation positions are given using the hg19 genome build. Depth, deduplicated tumour sequencing depth; panel_number, patient-specific mutation list number (multiple patients were grouped for panel design and analysis).

**Table S2** Sample library preparation input, QC, and INVAR likelihood ratios – test samples. For all patient samples, QC metrics, ctDNA IMAF values and sequencing depths are listed. Samples were non-evaluable if they had no ctDNA signal and <20,000 IR. SLX_barcode, unique identifier of the sample; DP_pre-dedup, mean depth per patient before deduplication;Study, study the sample belongs to; sample_type, the type of the sample used in the experiment; case_or_control, splitting samples into cancer patients and healthy controls; sequencer, sequencing platform the sample was sequenced on; library_prep, library preparation method used; input_into_library_ng, ng used for library preparation; patient, patient ID of the sample; Timepoint, timepoint the sample was taken; data_type, custom capture, WES or sWGS; IMAF, integrated mutant allele fraction; INVAR_LR, likelihood ratio; IR, informative reads; mut_sum, total sum of mutant collapsed reads observed in the sample; detected_using_size, sample was detected positive for ctDNA using size weighting approach; cancer_genomes_fraction, indicates the number of cancer genomes present in the sample, based on the number of mutant reads and the number of loci targeted; targeted_mutations, number of mutated loci targeted in the sample; hGA, haploid genomes analysed; LS_PASS, indicating samples passing low sensitivity threshold (set at > 20,000IR). This filter is not applicable to the whole genome sequencing cohort; n_timepoints, number of timepoints that were merged for this analysis.

**Table S3** Sample library preparation input, QC, and INVAR likelihood ratios – control samples. For all control samples, QC metrics, ctDNA IMAF values and sequencing depths are listed. SLX_barcode, unique identifier of the sample; DP_pre-dedup, mean depth per patient before deduplication;Study, study the sample belongs to; sample_type, the type of the sample used in the experiment; case_or_control, splitting samples into cancer patients and healthy controls; sequencer, sequencing platform the sample was sequenced on; library_prep, library preparation method used; input_into_library_ng, ng used for library preparation; individual, sample ID of the analyzed sample; data_type, custom capture, WES or sWGS; patient_loci_belong_to control samples are applied to mutation lists of patients on the same panel; IMAF, integrated mutant allele fraction; INVAR_LR, likelihood ratio; IR, informative reads; mut_sum, total sum of mutant collapsed reads observed in the sample; detected_using_size, sample was detected positive for ctDNA using size weighting approach; targeted_mutations, number of mutated loci targeted in the sample; cancer_genomes_fraction, indicates the number of cancer genomes present in the sample, based on the number of mutant reads and the number of loci targeted; hGA, haploid genomes analysed.

**Table S4** INVAR score thresholds. This table gives details on each of the cohorts, the experimental method performed to generate data, and the INVAR score threshold used (determined by ROC analysis). LR_threshold, likelihood ratio threshold used for detection; analytical_specificity, determined using other patients as control samples;

clinical_specificity, determined using healthy individuals on the same panel; min_family_size, minimum family size setting was used in read-collapsing.

**Table S5** Tumor volumes for stage IV melanoma cohort. This table shows the CT imaging data for stage IV melanoma patients. Days, days since start of treatment; n_lesions, number of lesions in the patient; evaluable, denotes evaluability of lesion through CT imaging. NA indicates time points where CT imaging was performed where there was not a corresponding plasma sample analysed.

**Table S6** Patient baseline characteristics. Baseline characteristics for each of the patient cohorts in this study are shown.

### Supplementary References

1. J. Zhang, K. Kobert, T. Flouri, A. Stamatakis, PEAR: A fast and accurate Illumina Paired-End reAd mergeR, *Bioinformatics* **30**, 614–620 (2014).
2. J. Phallen, M. Sausen, V. Adleff, A. Leal, C. Hruban, J. White, V. Anagnostou, J. Fiksel, S. Cristiano, E. Papp, S. Speir, T. Reinert, M.-B. W. Orntoft, B. D. Woodward, D. Murphy, S. Parpart-Li, D. Riley, M. Nesselbush, N. Sengamalay, A. Georgiadis, Q. K. Li, M. R. Madsen, F. V. Mortensen, J. Huiskens, C. Punt, N. van Grieken, R. Fijneman, G. Meijer, H. Husain, R. B. Scharpf, L. A. Diaz, S. Jones, S. Angiuoli, T. Ørntoft, H. J. Nielsen, C. L. Andersen, V. E. Velculescu, Direct detection of early-stage cancers using circulating tumor DNA, *Sci. Transl. Med.* **9** (2017) (available at http://stm.sciencemag.org/content/9/403/eaan2415.abstract).
3. C. Abbosh, N. J. Birkbak, G. A. Wilson, M. Jamal-Hanjani, T. Constantin, R. Salari, J. Le Quesne, D. A. Moore, S. Veeriah, R. Rosenthal, T. Marafioti, E. Kirkizlar, T. B. K. Watkins, N. McGranahan, S. Ward, L. Martinson, J. Riley, F. Fraioli, M. Al Bakir, E. GrÖnroos, F. Zambrana, R. Endozo, W. L. Bi, F. M. Fennessy, N. Sponer, D. Johnson, J. Laycock, S. Shafi, J. Czyzewska-Khan, A. Rowan, T. Chambers, N. Matthews, S. Turajlic, C. Hiley, S. M. Lee, M. D. Forster, T. Ahmad, M. Falzon, E. Borg, D. Lawrence, M. Hayward, S. Kolvekar, N. Panagiotopoulos, S. M. Janes, R. Thakrar, A. Ahmed, F. Blackhall, Y. Summers, D. Hafez, A. Naik, A. Ganguly, S. Kareht, R. Shah, L. Joseph, A. M. Quinn, P. Crosbie, B. Naidu, G. Middleton, G. Langman, S. Trotter, M. Nicolson, H. Remmen, K. Kerr, M. Chetty, L. Gomersall, D. A. Fennell, A. Nakas, S. Rathinam, G. Anand, S. Khan, P. Russell, V. Ezhil, B. Ismail, M. Irvin-sellers, V. Prakash, J. F. Lester, M. Kornaszewska, R. Attanoos, H. Adams, H. Davies, D. Oukrif, A. U. Akarca, J. A. Hartley, H. L. Lowe, S. Lock, N. Iles, H. Bell, Y. Ngai, G. Elgar, Z. Szallasi, R. F. Schwarz, J. Herrero, A. Stewart, S. A. Quezada, P. Van Loo, C. Dive, C. J. Lin, M. Rabinowitz, H. J. Aerts, A. Hackshaw, J. A. Shaw, B. G. Zimmermann, C. Swanton, M. Jamal-Hanjani, C. Abbosh, S. Veeriah, S. Shafi, J. Czyzewska-Khan, D. Johnson, J. Laycock, L. Bosshard-Carter, G. Goh, R. Rosenthal, P. Gorman, N. Murugaesu, R. E. Hynds, G. Wilson, N. J. Birkbak, T. B. K. Watkins, N. McGranahan, S. Horswell, M. Al Bakir, E. GrÖnroos, R. Mitter, M. Escudero, A. Stewart, P. Van Loo, A. Rowan, H. Xu, S. Turajlic, C. Hiley, J. Goldman, R. K. Stone, T. Denner, N. Matthews, G. Elgar, S. Ward, J. Biggs, M. Costa, S. Begum, B. Phillimore, T. Chambers, E. Nye, S. Graca, M. Al Bakir, K. Joshi, A. Furness, A. Ben Aissa, Y. N. S. Wong, A. Georgiou, S. Quezada, J. A. Hartley, H. L. Lowe, J. Herrero, D. Lawrence, M. Hayward, N. Panagiotopoulos, S. Kolvekar, M. Falzon, E. Borg, T. Marafioti, C. Simeon, G. Hector, A. Smith, M. Aranda, M. Novelli, D. Oukrif, A. U. Akarca, S. M. Janes, R. Thakrar, M. Forster, T. Ahmad, S. M. Lee, D. Papadatos-Pastos, D. Carnell, R. Mendes, J. George, N. Navani, A. Ahmed, M. Taylor, J. Choudhary, Y. Summers, R. Califano, P. Taylor, R. Shah, P. Krysiak,

K. Rammohan, E. Fontaine, R. Booton, M. Evison, P. Crosbie, S. Moss, F. Idries, L. Joseph, P. Bishop, A. Chaturved, A. Marie Quinn, H. Doran, A. Leek, P. Harrison, K. Moore, R. Waddington, J. Novasio, F. Blackhall, J. Rogan, E. Smith, C. Dive, J. Tugwood, G. Brady, D. G. Rothwell, F. Chemi, J. Pierce, S. Gulati, B. Naidu, G. Langman, S. Trotter, M. Bellamy, H. Bancroft, A. Kerr, S. Kadiri, J. Webb, G. Middleton, M. Djearaman, D. Fennell, J. A. Shaw, J. Le Quesne, D. Moore, A. Thomas, H. Walter, J. Riley, L. Martinson, A. Nakas, S. Rathinam, W. Monteiro, H. Marshall, L. Nelson, J. Bennett, J. Riley, L. Primrose, L. Martinson, G. Anand, S. Khan, A. Amadi, M. Nicolson, K. Kerr, S. Palmer, H. Remmen, J. Miller, K. Buchan, M. Chetty, L. Gomersall, J. Lester, A. Edwards, F. Morgan, H. Adams, H. Davies, M. Kornaszewska, R. Attanoos, S. Lock, A. Verjee, M. MacKenzie, M. Wilcox, H. Bell, N. Iles, A. Hackshaw, Y. Ngai, S. Smith, N. Gower, C. Ottensmeier, S. Chee, B. Johnson, A. Alzetani, E. Shaw, E. Lim, P. De Sousa, M. Tavares Barbosa, A. Bowman, S. Jordan, A. Rice, H. Raubenheimer, C. Proli, M. Elena Cufari, J. C. Ronquillo, A. Kwayie, H. Bhayani, M. Hamilton, Y. Bakar, N. Mensah, L. Ambrose, A. Devaraj, S. Buderi, J. Finch, L. Azcarate, H. Chavan, S. Green, H. Mashinga, A. G. Nicholson, K. Lau, M. Sheaff, P. Schmid, J. Conibear, V. Ezhil, B. Ismail, M. Irvin-sellers, V. Prakash, P. Russell, T. Light, T. Horey, S. Danson, J. Bury, J. Edwards, J. Hill, S. Matthews, Y. Kitsanta, K. Suvarna, P. Fisher, A. D. Keerio, M. Shackcloth, J. Gosney, P. Postmus, S. Feeney, J. Asante-Siaw, T. Constatin, R. Salari, N. Sponer, A. Naik, B. Zimmermann, M. Rabinowitz, H. J. W. L. Aerts, S. Dentro, C. Dessimoz, C. Swanton, M. Jamal-Hanjani, C. Abbosh, K.-K. Shiu, J. Bridgewater, D. Hochauser, P. Van Loo, S. Quezada, S. Beck, P. Parker, H. Walczak, T. Enver, M. Falzon, I. Proctor, R. Sinclair, C. Lok, M. Novelli, T. Marafioti, E. Borg, M. Mitchison, G. Trevisan, M. Lynch, S. Brandner, F. Gishen, A. Tookman, P. Stone, C. Sterling, J. Larkin, S. Turajlic, G. Attard, R. Eeles, C. Foster, S. Bova, A. Sottoriva, S. Chowdhury, C. Ashish, J. Spicer, M. Stares, J. Lynch, C. Caldas, J. Brenton, R. Fitzgerald, M. Jimenez-Linan, E. Provenzano, A. Cluroe, G. Stewart, C. Watts, R. Gilbertson, U. McDermott, S. Tavare, T. Maughan, I. Tomlinson, P. Campbell, I. McNeish, A. Biankin, A. Chambers, S. Fraser, K. Oien, M. Krebs, F. Blackhall, Y. Summers, C. Dive, R. Marais, L. Carter, D. Nonaka, A. M. Quinn, N. Dhomen, D. Fennell, J. Le Quesne, D. Moore, J. Shaw, B. Naidu, S. Baijal, B. Tanchel, G. Langman, M. Collard, P. Cockcroft, J. Taylor, H. Bancroft, A. Kerr, G. Middleton, J. Webb, S. Kadiri, P. Colloby, B. Olisemeke, R. Wilson, C. Ottensmeier, D. Harrison, M. Loda, A. Flanagan, M. Wilcox, M. McKenzie, A. Hackshaw, J. Lederman, A. Sharp, L. Farrelly, C. Swanton, Phylogenetic ctDNA analysis depicts early stage lung cancer evolution, *Nature* **22364**, 1–25 (2017).

4. H. C. Fan, Y. J. Blumenfeld, U. Chitkara, L. Hudgins, S. R. Quake, Analysis of the size distributions of fetal and maternal cell-free DNA by paired-end sequencing, *Clin. Chem.* **56**, 1279–1286 (2010).

5. M. López-Ratón, M. X. Rodríguez-Álvarez, C. C. Suárez, F. G. Sampedro, OptimalCutpoints : An R Package for Selecting Optimal Cutpoints in Diagnostic Tests, *J. Stat. Softw.* **61**, 1–36 (2014).

6. A. M. Newman, S V Bratman, J. To, J. F. Wynne, N. C. W. Eclov, L. a Modlin, C. L. Liu, J. W. Neal, H. a Wakelee, R. E. Merritt, J. B. Shrager, B. W. Loo, A. a Alizadeh, M. Diehn, An ultrasensitive method for quantitating circulating tumor DNA with broad patient coverage., *Nat. Med.* **20**, 548–54 (2014).
